# Prophage taming by the adherent-invasive *Escherichia coli* LF82 upon macrophage infection

**DOI:** 10.1101/2022.10.28.514194

**Authors:** Pauline Misson, Emma Bruder, Jeffrey K. Cornuault, Marianne De Paepe, Gaëlle Demarre, Marie-Agnès Petit, Olivier Espeli, François Lecointe

## Abstract

Adherent-invasive *Escherichia coli* (AIEC) strains are frequently recovered from stools of patients with dysbiotic microbiota. They have remarkable properties of adherence to the intestinal epithelium, and survive better than other *E. coli* in macrophages. The best studied of these AIEC is probably strain LF82, which was isolated from a Crohn’s disease patient. This strain contains five complete prophages, which have not been studied until now. We undertook their analysis, both *in vitro* and inside macrophages, and show that all of them form virions. The Gally prophage is by far the most active, generating spontaneously over 10^8^ viral particles per mL of culture supernatants *in vitro*, more than 100-fold higher than the other phages. Gally is over-induced after a genotoxic stress generated by ciprofloxacin and trimethoprim. However, upon macrophage infection, Gally virion production is decreased by more than 20-fold, and the transcription profile of the prophage indicates that part of the structural module is specifically repressed while the replication module is overexpressed compared to unstressed culture conditions. We conclude that strain LF82 has evolved an efficient way to “tame” its most active prophage upon macrophage infection, which may participate to its good survival in macrophages. The results are discussed in light of the active lysogeny process.

**AUTHOR SUMMARY:** Prophages are bacterial viruses stably integrated into their host, to which they can provide new functions, thus increasing their fitness in the environment. Thereby, they can participate to the virulence of bacterial pathogens. However, prophages are double-edged swords that can be awakened in response to genotoxic stresses, resulting in the death of their bacterial host. This raises the question of the effect of this type of stress in the natural environments where their bacterial hosts exert their virulence. In this study, we characterized the five active prophages present in *Escherichia coli* LF82, a strain belonging to the intestinal microbiota and suspected to be involved in Crohn’s disease via its ability to invade macrophages, a highly genotoxic environment. We show that LF82 inhibits the awakening of its prophages in macrophages, allowing it to survive there. Moreover, deletion of its most active prophage does not affect the viability of LF82 in this environment. These results show that LF82 has tamed its prophages in macrophages and also suggest that if they convey fitness advantages, they probably do so in environments differing from macrophages, and which remain to be discovered.

## INTRODUCTION

Lysogens, the bacteria hosting functional prophages (either integrated into their genome or as freely replicating episomes), are abundant in natural ecosystems, representing approximately half of all completely sequenced strains (1), and up to 70% of strains in species of the human microbiota (2–5). Lysogeny is often considered as beneficial for the bacterium, thanks to the expression of prophage genes named “morons”. Morons are genes that are autonomously expressed (i.e. not under the control of the lysogeny master regulator), are not involved in the lytic cycle of the phage but provide the bacterial host with some gain of function (for a review see (6)). Moron functions are diverse, ranging from protection against infection by other phages (7–9), metabolic genes (2, 10) or adaptation to a given environment (11, 12), and many remain to be characterized. However, this potential benefit of lysogeny is counterbalanced by the permanent danger of lysis, due to prophage induction, i.e. its entry into a lytic cycle upon derepression of the lysogeny master regulator. Bacterial growth may be hindered for lysogens compared to non lysogens if the prophage is constantly induced in a significant proportion of the population. This burden is important in the case of an *Escherichia coli* lysogen propagated in the gastro-intestinal tract (GIT) of monoxenic mice (13), and has also been observed for *Lactobacillus reuterii* lysogens, during transit in conventional mice (3). In both cases, prophage induction was RecA-dependent, suggesting that the inducing signal was due to genotoxic stress. In the mammalian gut, molecules activating the SOS response, might not only be produced by the host itself, but also by the microbiota (14): for *L. reuterii*, fructose consumption and acetic acid release was suggested as the origin of the genotoxic stress inducing its prophages in the GIT (3). Few RecA-independent pathways have also been described for prophage induction (15–18) and other sensors and pathways certainly await discovery.

Characterizing lysogeny, and the signals regulating prophage induction in natural settings, is therefore critical for the understanding of bacterial behaviors in their natural environments, and more particularly those of bacterial pathogens. Indeed, bacterial pathogens are often lysogens, and even poly-lysogens (1). The first prophage morons to be described were virulence factors, such as the shiga-toxin, to mention just one of many important phage-encoded toxins (6). All happens as if prophages were adjustment variables, allowing pathogens to rapidly adapt to new niches and compete with the local inhabitants (19). The adherent-invasive *Escherichia coli* (AIEC) genomes are also richly decorated with prophages, and strain LF82, its best characterized member, encodes five prophages predicted to be complete and functional, based on genomic analyses (20, 21). One of these, named “prophage 1” was even reported as significantly associated to the *E. coli* strains isolated from Crohn’s disease patients (22). Yet, whether these prophages are functional (able to complete lytic cycles, form virions and multiply), and what kind of signal or stress induces them, had not been investigated.

AIEC have two remarkable properties: they adhere to the intestinal epithelium (23), and they invade and multiply in macrophages (24, 25), being able to form biofilm within vacuoles (26).

Interestingly, it was recently established that the macrophage environment can provoke Lambda prophage induction by 26-fold compared to *in vitro* in an unstimulated model laboratory *E. coli* strain (15). This high induction level, which leads *in vitro* to the lysis of ∼90% of the bacteria, probably facilitates the clearing work of macrophages (15). In the vacuolar compartment, Lambda induction was not dependent on RecA, as observed *in vitro*, but rather on PhoP. An anti-microbial peptide, mCramp1, was suggested to be at the origin of this induction, via a bacterial membrane stress (15). Whether prophages from natural *E. coli* isolates are also induced and provoke bacterial lysis in macrophages remains unknown. This question is of importance for LF82 that survives in macrophages and contains five putative active prophages.

To further understand the good survival of LF82 in macrophages, we undertook the characterization of its prophages, both *in vitro* and inside macrophages. We show that the five prophages form virions *in vitro*. One of them, the phage Gally (formerly prophage 1) dominates the culture supernatants: its spontaneous induction level is high enough to generate above 10^8^ particles per mL of culture during exponential growth in rich medium. Interestingly, and contrary to expectations, production of Gally virions was reduced over 20-fold upon growth in macrophages. We hypothesize that the remarkable survival of LF82 in macrophage is partly due to its ability to control the induction level of its most active prophage.

## RESULTS

### The prophages of LF82 encode various morons

Previous analysis of the *E. coli* LF82 genome had identified four putative complete prophages and one plasmid, pLF82, subsequently found to be homologous to the *Salmonella* SSU5 phage (20). We named them here Gally (previously prophage 1), Perceval (prophage 2), Tritos (prophage 3), Cartapus (prophage 4) and Cyrano (pLF82), and upon updated re-annotation, these were introduced into the European Nucleotide Archive database (see Material and Methods). The phage-plasmid Cyrano closely resembles phage SSU5 (81% identity, 65% coverage, see S1 Fig), and is predicted to have a siphovirus morphology. Among integrated prophages, Gally is partially homologous to the podophage HK620 (97% identity on 44% coverage, Fig 1 top), and follows the general genetic organization of P22 (Lederbergvirus genus). Perceval is a close relative of Ev207, a phage isolated from the infant gut and homologous to Lambda (2). Tritos is also distantly related to Lambda, and close to another infant gut phage, named Ev081 (2). Finally, Cartapus is a P2- related phage, closely similar to Fels2 (Felsduovirus genus).

**Fig 1.**
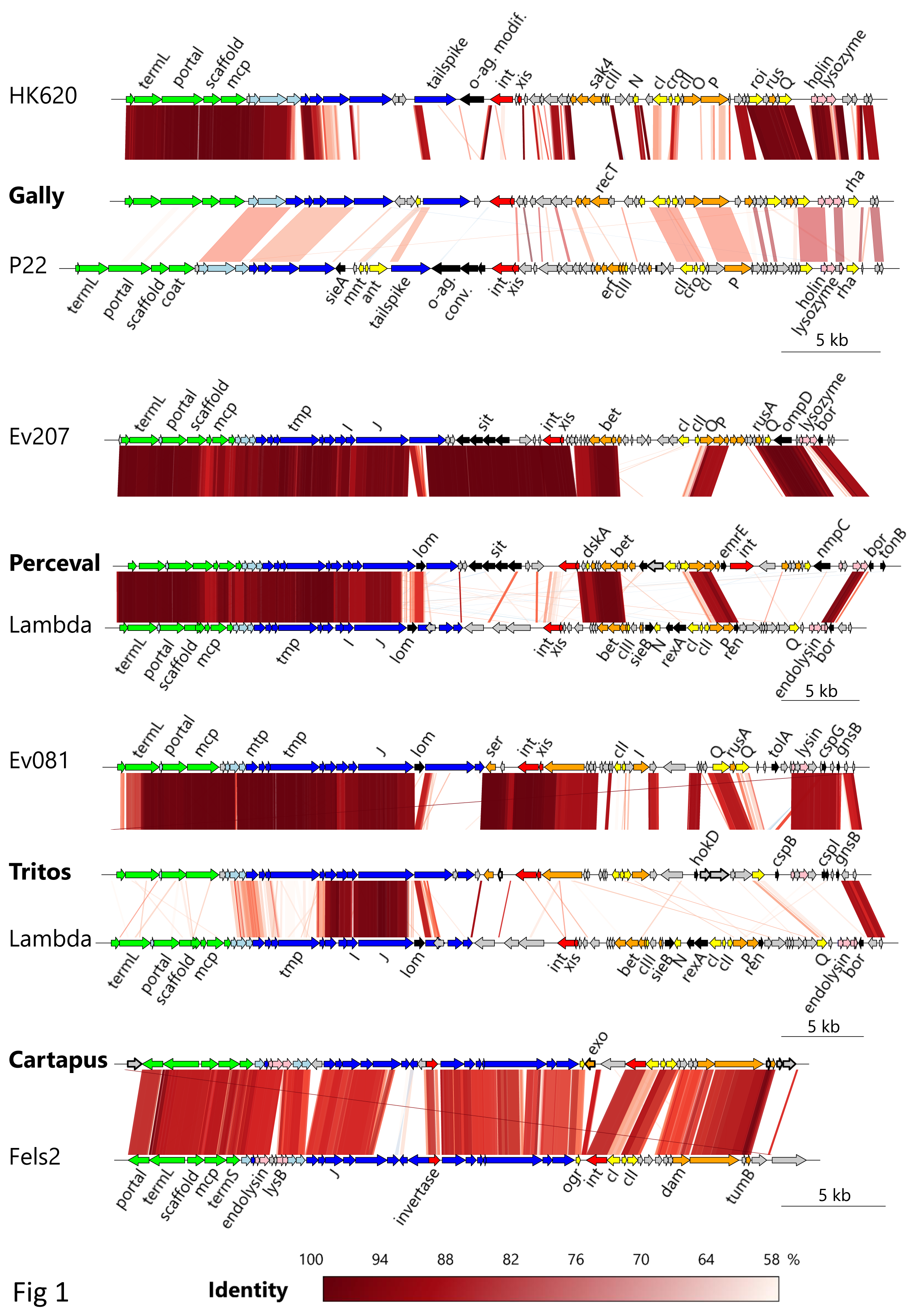
Whole genome comparisons of the four integrated prophages. Each LF82 phage (in bold) is compared with tBLASTx to its closest relative in databases (up), as well as to an ICTV classified prototype (down), and displayed using the R Genoplot package. Gene color code: green, capsid; light blue, connector; dark blue, tail; red, integration and excision; orange, DNA metabolism; yellow, transcriptional regulators; pink, lysis; grey, hypothetical. Morons displaying sequence homology with predicted or validated known morons are indicated as black arrows whereas morons newly identified by our transcriptomic analysis (see Material and methods) are indicated as grey arrows with thick black border.

To identify the morons of the five prophages, transcriptomes of LF82 grown *in vitro* either in LB (2 repeats) or in NMDM (a rich medium for macrophage propagation, 2 repeats) were analyzed (26). Morons were identified as genes (i) different from the master regulator or other typical phage genes (ii) pertaining to autonomous transcription units, and (iii) transcribed at least 5-fold above the local transcriptional background of the prophage region (see Material and Methods, Fig 1). Cyrano and Perceval were the richest in moron content (10 and 8 genes), followed by Tritos and Cartapus (7 and 5 genes) while Gally was apparently devoid of any moron (Table 1). Among the morons with predicted functions (18 of the 30), Perceval encoded an iron and manganese transporter SitABCD and an EmrE multidrug transporter.

**Table 1:** Prophage genes expressed at least 5-fold above local background in strain LF82 grown in LB medium (overnight culture) or in DMEM (exponentially for 1h).

| *Prophage name* | *Locus tag* | *Annotation* | *Fold above Background<sup>a</sup>* |
| --- | --- | --- | --- |
| *Gally* | *LF82_107* | *Repressor protein<sup>b</sup>* | 69 |
| *Perceval* | *LF82_154* | *cI repressor* | 38 |
| Perceval | LF82_152 | hypothetical protein | 128 |
| Perceval | LF82_153 | hypothetical protein | 48 |
| Perceval | LF82_0556 | EmrE Multidrug transporter | 16 |
| Perceval | LF82_160 | TonB membrane protein, partial | 10 |
| Perceval | LF82_184 | SitD iron transport subunit | 20 |
| Perceval | LF82_185 | SitC iron transport subunit | 20 |
| Perceval | LF82_186 | SitB iron transport subunit | 39 |
| Perceval | LF82_187 | SitA iron transport subunit | 63 |
| *Tritos* | *LF82_287* | *cI repressor* | 37 |
| Tritos | LF82_2833 | YdfK Cold shock protein | 7 |
| Tritos | LF82_0907 | GnsB toxin | 43 |
| Tritos | LF82_0377 | CspI cold shock protein | 54 |
| Tritos | LF82_0371 | CspB Cold shock-like protein | 147 |
| Tritos | LF82_280 | ImmA/IrrE metallo-endopeptidase | 35 |
| Tritos | LF82_281 | hypothetical protein | 53 |
| Tritos | LF82_1023 | Hok toxin | 3272 |
| *Cartapus* | *LF82_791* | *cI repressor* | 83 |
| Cartapus | LF82_783 | putative exonuclease protein | 168 |
| Cartapus | LF82_788 | hypothetical protein | 37 |
| Cartapus | LF82_789 | hypothetical protein | 44 |
| Cartapus | LF82_790 | hypothetical protein | 13 |
| Cartapus | LF82_791 | hypothetical protein | 20 |
| *Cyrano* | *CAP72145.1* | *Replication protein RepA* | 14 |
| Cyrano | CAP72157.1 | HicA toxin protein | 95 |
| Cyrano | CAP72158.1 | putative HicB antitoxin | 43 |
| *Cyrano* | *CAP72163.1* | *cI repressor* | 26 |
| Cyrano | CAP72165.1 | hypothetical protein | 602 |
| Cyrano | CAP72177.1 | hypothetical protein | 22 |
| Cyrano | CAP72178.1 | RelE/ParE putative toxin | 44 |
| Cyrano | *CAP72179.1* | *ParA plasmid partitioning protein* | 22 |
| Cyrano | CAP72180.1 | putative antitoxin | 109 |
| Cyrano | CAP72181.1 | KacT Acetyltransferase-type toxin | 70 |
| Cyrano | CAP72213.1 | hypothetical protein | 37 |
| Cyrano | CAP72221.1 | hypothetical protein | 7 |
| Cyrano | *CAP72232.1* | *LexA-like peptidase S24* | 37 |
| Cyrano | CAP72249.1 | hypothetical protein | 27 |

<sup>a</sup> Fold over background corresponds to the ratio of the mean RPKM value for each phage gene on the median value of RPKM of all genes from the same prophage in these two growth conditions (see Materials and Methods). <sup>b</sup> Master repressors of lysogeny, as well as the few other expressed genes of the phage-plasmid Cyrano, are shown in italics, as a reference.

The Bor transcript, an early recognized moron and outer membrane protein involved in serum resistance (12), was actively transcribed in LB (50-fold above background) but not in NMDM (3- fold above background), explaining its absence in Table 1. Tritos and Cyrano encoded several toxins (GnsB and HokD for Tritos, KacT, RelE and HicA for Cyrano) as well as an annotated antitoxin (HicB) in Cyrano. Among these morons, one was also a recognized virulence factor: the *sitABCD* operon. We conclude that all prophages but Gally are rich in morons.

Inspection of attachment sites of the four integrated prophages did not reveal any particular gene inactivation pattern. Notably however, Gally was placed between the divergent *torS* (the sensor of the *torS/R* system, pointing leftward) and *torT* (a periplasmic protein pointing rightward). This site is the target of prophage insertion in approximately 5% of *E. coli* strains (27). The presence of the HK022 prophage at this site increases the expression of *torS* and consequently inhibits the expression of the *torCAD* operon when cells are grown aerobically (27). The *torCAD* operon codes for a trimethylamine N-oxide (TMAO) reductase that allows the respiration of TMAO by *E. coli* (28).

### The five LF82 prophages are spontaneously induced *in vitro*

We next asked whether some or all of LF82 prophages were producing virions. To identify prophages spontaneously producing viral particles, we filter purified the virome from the supernatant of an LF82 culture grown in rich medium until stationary phase, treated it with DNase I before destroying capsids, and sequenced the encapsidated DNA. The vast majority of reads (94.6 %, mean coverage= 35,319 reads/bp), mapped on Gally (Fig 2), indicating that this phage is highly produced from LF82 cultures. The next most induced prophage was Tritos (0.14% of the mapped reads, mean coverage 24.6, compared to 2.5 in the 2 kb flanking regions) and to a lesser extent Cartapus (0.03% of the mapped reads, mean coverage = 13.7 compared to 7.6 around). The activity of Perceval could not be evaluated by this method, due to its proximity with Gally on the LF82 chromosome (180 kb), combined with the ”leakage” of the Gally signal over the Perceval region (see the lateral transduction section below). Finally, a low noise of reads mapped on the rest of LF82 genome (4.3% of total, mean coverage =7.6), corresponding either to contaminating or transduced bacterial DNA.

**Fig 2.**
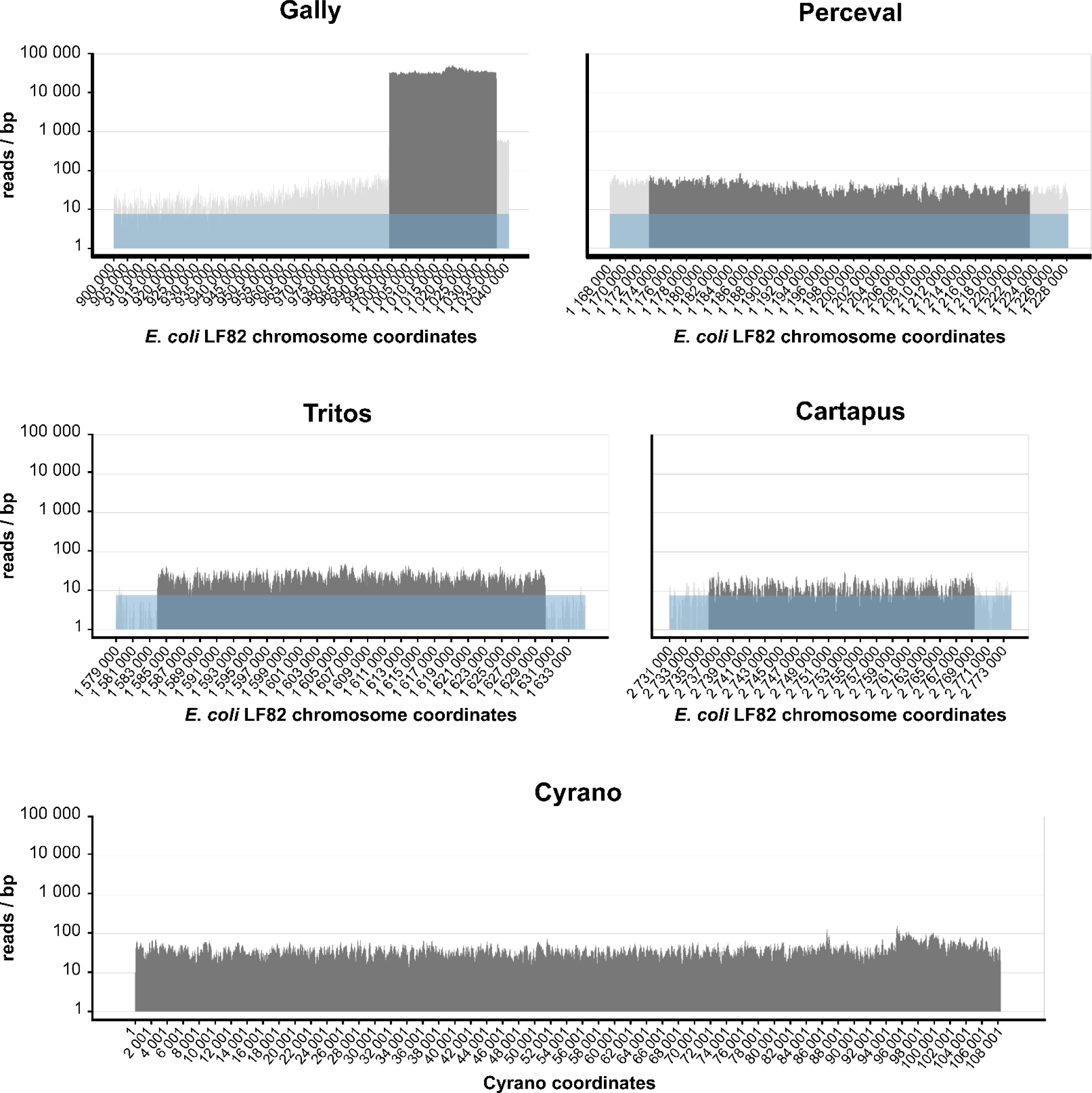
Shotgun sequencing of the viral particles generated by the five LF82 prophages in an *in vitro* culture in rich medium. Nucleotide coverage as a function of LF82 chromosome or Cyrano phage-plasmid coordinates obtained after the mapping of the sequencing reads from the virome of an overnight LF82 culture. The coverages associated with the prophage regions and the bacterial chromosome are shown in dark and light grey respectively. The average coverage linked to bacterial DNA contamination of the virome sequencing is represented in light blue (7.6 reads/bp).

The mean coverage of the phage-plasmid Cyrano was 41.2 (0.74% of total reads). We took advantage of published Tn-seq analysis (26) to estimate the copy number of this phage-plasmid. We observed an average 3-fold increase of transposon insertions in Cyrano compared to the chromosome, suggesting a copy number of 3 for Cyrano. The mean coverage of Cyrano was therefore 1.8-fold above its expected value of 22.8 (3 * 7.6), suggesting a mild induction level in LF82 cultured in unstressed conditions.

To confirm the prevalence of Gally in culture supernatants, virions were quantified by quantitative PCR (qPCR), on ten independent filtered culture supernatants, grown to exponential phase in unstressed conditions. As a control, a qPCR on the *ybtE* gene of LF82, coding for the yersiniabactin biosynthesis salycil-AMP ligase, allowed to estimate a mean value of bacterial contamination for the virome preparations of 2.3x10^4^ chromosomal copies/mL. As expected from the virome sequencing, a high concentration of 4.3x10^8^ Gally genomes per mL was measured (Fig 3, no antibiotic). Knowing that the bacterial concentration was approximately 5x10^8^ CFU/mL, the ratio of Gally virions per bacteria was around 1 in this unstressed growth condition. qPCR were also performed for the other phages and revealed concentrations two to three orders of magnitude below Gally for Cyrano, Tritos and Cartapus, which produced 4.2x10^6^, 2.8x10^6^ and 3.3x10^5^ genomes per mL respectively. Due to the lateral transduction of Gally (see below), qPCR using primer targeting Perceval DNA allowed to measure 1.5x10^6^ virions per mL that contain the whole or only a part of the Perceval genome.

**Fig 3.**
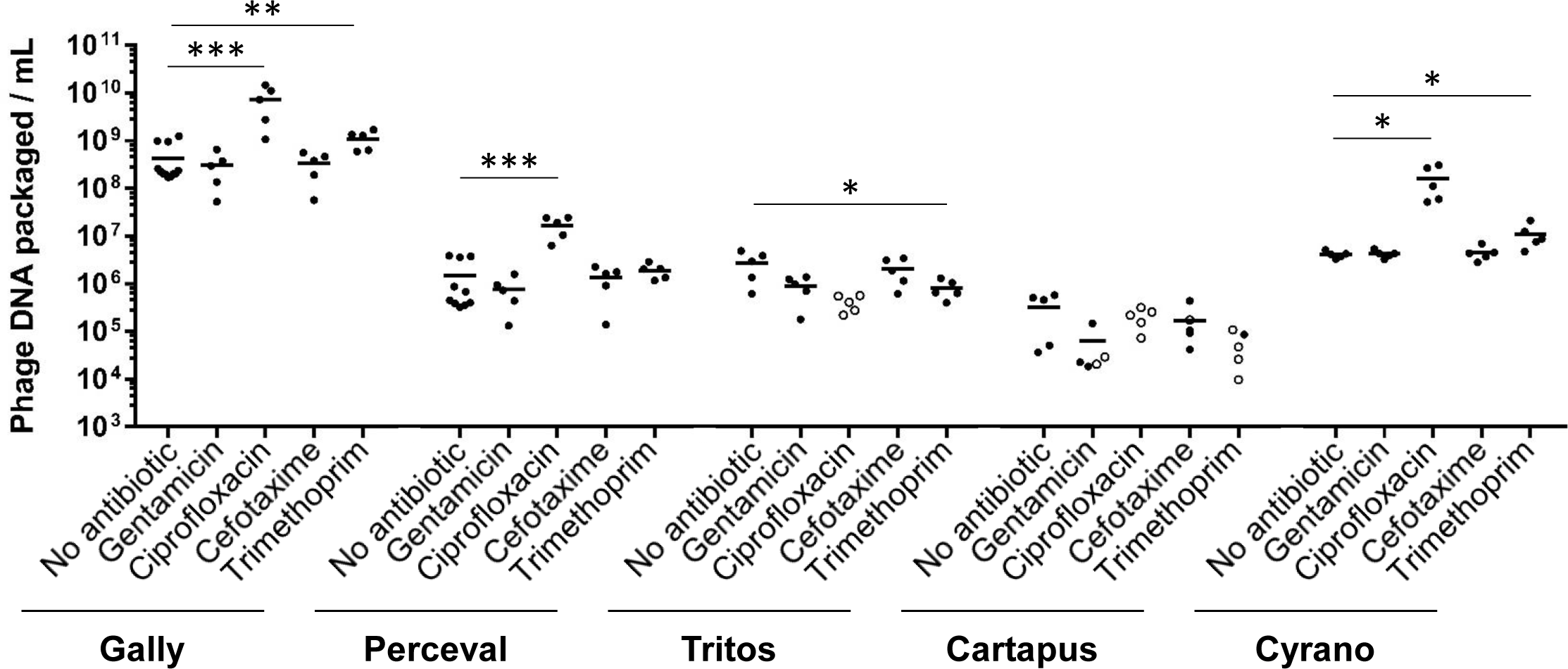
PCR quantification of the viral particles generated by the five LF82 prophages in rich medium with or without antibiotic from an *in vitro* LF82 culture. Phages produced from LF82 cells cultured in presence or absence of an antibiotic, as indicated, were quantified. Each dot corresponds to one biological replicate. Black or white dots correspond to replicates that are more or less abundant than their associated LF82 genomic DNA contamination respectively. Only values corresponding to black dots are used to calculate the mean (vertical line). The statistical difference (t-test) between antibiotic treated and untreated cultures is indicated by one (pvalue < 0.05), two (pvalue < 0.1) or three (pvalue < 0.005) asterisks.

We next attempted to propagate and purify these phages as plaques on an indicator strain, to allow their visualization by transmission electron microscopy (TEM, Fig 4). We succeeded in the isolation and visualization of the two siphoviruses Perceval (capsid diameter (c.d.): 63.3±1.6 nm, tail length (t.l.): 156.8±5.3 nm and tail thickness (t.t.): 11.3±1.5 nm) and Tritos (c.d.:57.2±9.6 nm, t.l.: 163.5±2.4 nm, t.t.: 11.3±0.4 nm). Despite its abundance, Gally could not be propagated under all conditions tested (see Material and Methods) but two Gally-Perceval hybrids (named Galper1 and Galper2, S2 Fig) were isolated instead. Galper1 displayed capsid and tail dimensions similar to those of Perceval (c.d.: 62.1±2.3nm, t.l.: 159.1±5.6 nm and t.t.: 12.7±2.1 nm), and its genome contained all structural and lysis genes from Perceval, interrupted by the replication module of Gally (S2 Fig). The junctions consisted in short homology regions (S2 Fig), typical of the substrates used by phage single strand annealing protein (SSAP) (29, 30). The rightward recombination junction was identical in Galper2, but the left one (region Rz) was slightly offset (S2 Fig).

**Fig 4.**
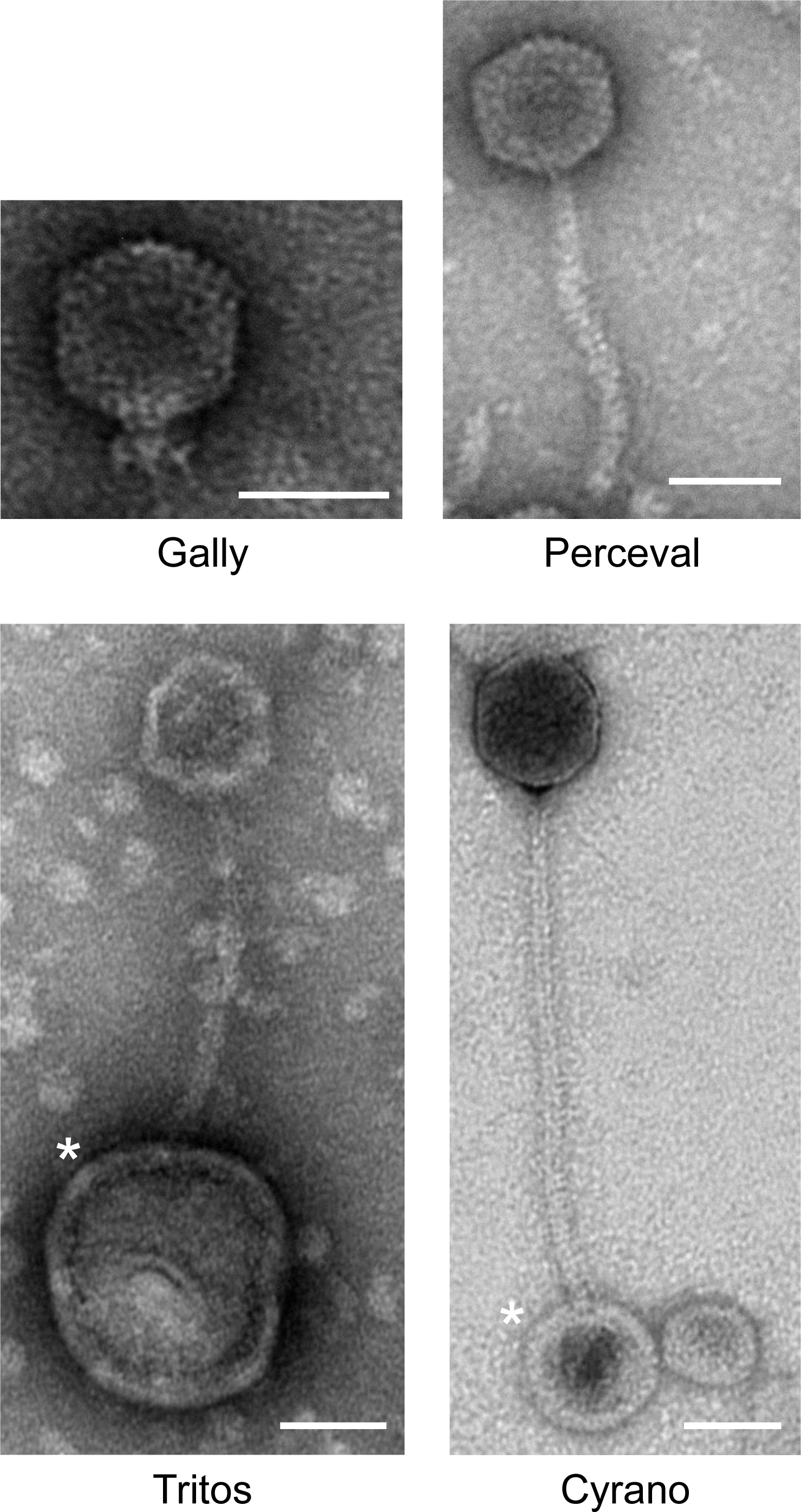
Transmission electron microscopy photographs of the different virions produced by LF82. Gally and Cyrano were imaged directly from LF82 culture supernatants. Perceval and Tritos phages were visualized after their purification and propagation on MAC1403 as an indicative strain. Asterisks indicate vesicles. Scale bars are 50 µm long.

In order to visualize Gally virions, we took advantage of its abundance in LF82 culture supernatants and its sequence homology with the HK620 and P22 podoviruses. TEM images of an overnight LF82 culture supernatant showed a huge abundance of a podovirus (c.d.: 69.2±1.9 nm), that we surmised corresponded to Gally (Fig 4). To search for Cyrano virions, we started from a ciprofloxacin treated LF82 culture (see below) and screened for siphoviruses displaying a tail around 228 nm in length (applying the 0.15 nm/ amino acids rule to the 1525 amino acids long TMP protein (31, 32)). Virions with a large head (c.d.: 76.4±4.0 nm capsid diameter) and a long non-contractile tail (t.l.: 259.1±11.9 nm, t.t.: 10.3±0.4 nm) were found. This virion being the only siphovirus remaining to identify among phages produced by LF82, and having dimensions typical for SSU5 phages, we hypothesized that it was Cyrano (Fig 4). Finally, no Cartapus-like myoviruses were visualized by TEM, as no recipient strain for its propagation was found, and its abundance was always low in viromes.

Overall, by combining virome isolation and sequencing, qPCR quantification and electronic microscopy on virome sample or phage isolates, we can conclude that under unstressed *in vitro* growth conditions, the five LF82 prophages are induced and form virions. Among them, Gally is by far the most abundant. Of note, for two phages only (Perceval and Tritos) out of the five, we identified sensitive hosts proving that these are also infectious.

### Important lateral transduction mediated by the *pac* type phage Gally

Further analysis of sequencing reads mapping on the LF82 chromosome revealed a particular property of phage Gally (Fig 5). A mean coverage of the Gally prophage of ∼35,000 was observed (dark grey region, Fig 5), corresponding to the bulk of encapsidated DNA. This coverage was not homogenous all along the prophage however, as a sharp peak was standing out, localized within the *termS* gene coding for the small terminase subunit (position 1,019,224 on LF82) and showing a gradual rightward decrease until the prophage *attR* attachment site (Fig 5A). Such a pattern is a signature of a headful packaging mechanism, initiated at the *pac* site, localized at the left end side of the peak (Fig 5A).

**Fig 5.**
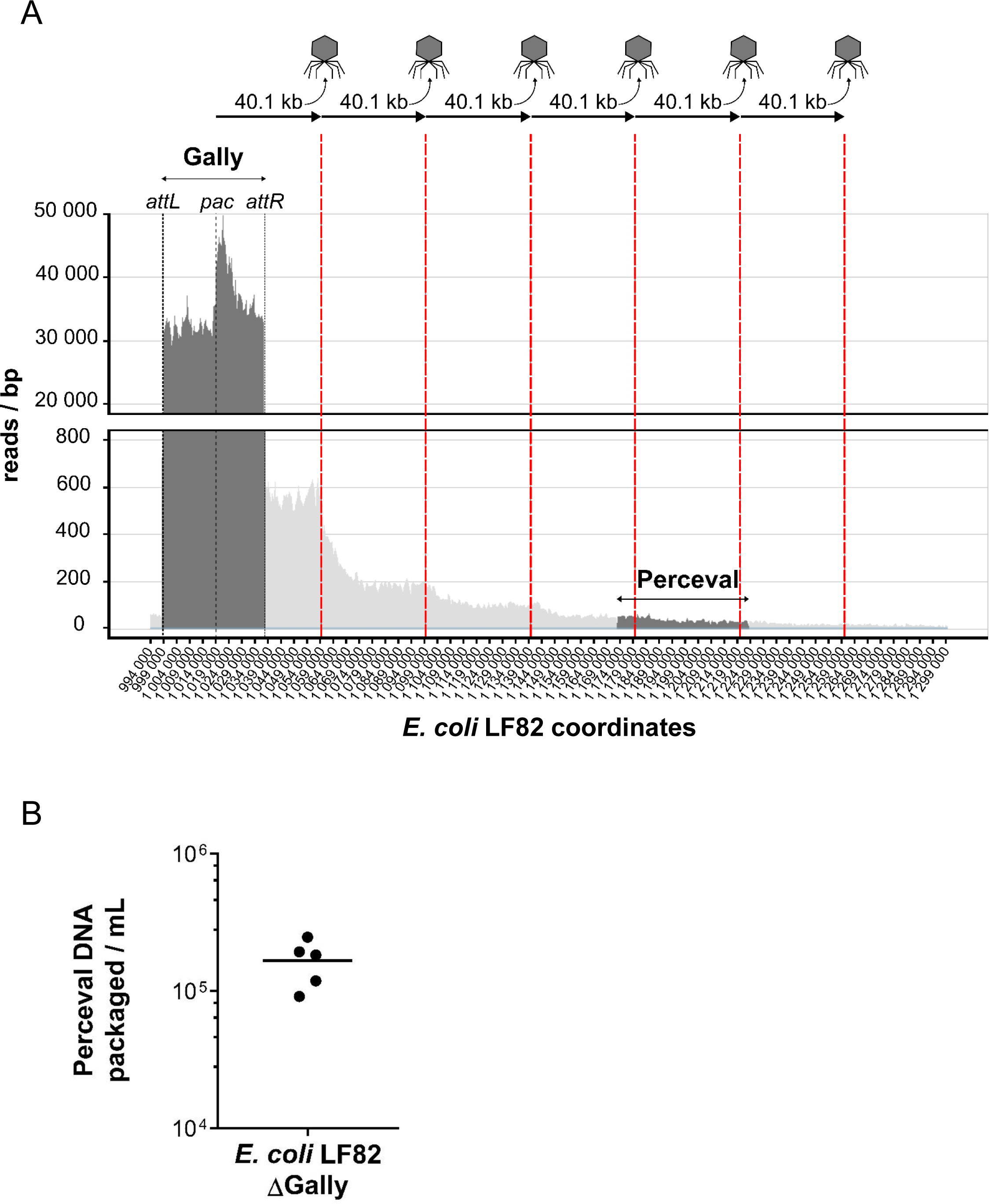
Gally performs lateral transduction of over 200 kb of adjacent chromosomal DNA, including Perceval. A. Zoom of mapped reads within the chromosomal region to the right of Gally prophage containing Perceval. The abrupt increase in reads inside Gally corresponds to the *pac* region, from which the packaging is initiated. Red bars indicate 40.1 kb steps of decreasing coverage from the *pac* site of Gally prophage, which could correspond to DNA packaged thanks to a headful mechanism (103.8% of the Gally genome length). B. PCR quantification of Perceval virions produced from an *in vitro* culture in reach medium of *E. coli* LF82 strain deleted for the Gally prophage. Each black dot corresponds to one biological replicate and the black horizontal line indicates the mean value.

Furthermore, adjacent to the rightward Gally *attR*, we observed, as mentioned above, a “leakage” where read coverage was well above background, and organized in successive steps of decreasing coverage values. The read coverage immediately downstream the *attR* site was 65-fold higher than the background (coverage ∼500 compared to 7.6) and was stable over ∼20 kb. Following this first reduction, the read coverage decreased about two to three fold every 40 kb on this side (Fig 5A). Considering that P22 packages ∼103.8% of its genome per capsid (33), Gally virions could contain ∼40.1 kb of DNA, which corresponds to the approximate length of each decreasing step in the read coverage. This profile of read coverage was described as a consequence of a specific transduction event processed by *pac*-type prophages, called lateral transduction that was initially described for *Staphylococcus* phage 80α (34), and then reported for P22 (35).

In line with the lateral transduction process, upstream of the *attL* site, the read coverage was approximately 7-fold higher than average DNA contamination (∼50 vs 7.6) and decreased, not by step as observed downstream the *attR* site, but progressively (Fig 2). This is likely the result of the *in situ* bi-directional replication of prophage Gally upon induction and before its delayed excision. As mentioned above, Perceval is covered by the lateral transduction area of Gally (Fig 5A), preventing the correct quantification of Perceval virions by DNA-based approaches. To investigate whether Perceval was an active prophage, we isolated a strain deleted of the Gally prophage (see Material and Methods). Culture of this mutant in unstressed growth condition led to the production of 1.7x10^5^ particles containing Perceval DNA / mL (Fig 5B). Comparison of the viral particles containing Perceval DNA produced by the wild-type strain (1.5x10^6^ particles containing Perceval DNA / mL, Fig 3) and the Gally prophage deleted strain indicated that approximately 90% of the particles containing Perceval DNA were the result of lateral transduction initiated by Gally in the wild-type LF82 strain, assuming a similar induction rate of Perceval in wild-type and ΔGally strains. Perceval is therefore the least produced virion by LF82 in this unstressed condition.

### At least two of the five prophages are induced by ciprofloxacin and trimethoprim

We next investigated whether some antibiotics could induce LF82 prophages beyond their spontaneous level. Genotoxic stresses are known to induce the lytic cycle of many phages *via* the activation of RecA and the cleavage of the master regulator of the lysogeny. We therefore tested antibiotics which activate RecA (and the downstream SOS response) to various extents: i) ciprofloxacin that inhibits DNA gyrase and topoisomerase IV activities leading to replication fork stalling, ii) trimethoprim which prevents synthesis of tetrahydrofolate leading subsequently to DNA damages and iii) cefotaxim, a beta-lactam antibiotics inhibiting primarily the peptidoglycan synthesis but also inducing the SOS response via inhibition of the replication (36, 37). We also tested gentamycin, an aminoglycoside that does not induce the SOS response, which we used to eliminate non-invading bacteria upon macrophage infection. The minimal inhibitory concentrations (MICs) for these antibiotics were first determined (see Material and Methods). Then, antibiotics were added at concentrations corresponding to the MIC in exponentially growing cultures, and supernatants were harvested two hours later, filtered and virions were quantified by qPCR. Gally and Cyrano were strongly induced by ciprofloxacin, 18 and 39-fold respectively (Fig 3). We could not conclude whether ciprofloxacin also induced Perceval, as the 12-fold increase in Perceval copy number was probably a consequence of the lateral transfer activity of Gally. Trimethoprim had a mild, if any, induction effect on Gally and Cyrano (2.7-fold), and decreased slightly (3.4-fold) the production of Tritos. Finally, cefotaxime and gentamycin did not have any effect on the induction level of the five prophages. We conclude that a genotoxic stress similar to the one provoked by a 2 hours ciprofloxacin-exposure at the MIC strongly induces part of the LF82 phageome.

### LF82 survival in macrophages is not affected by the presence of Gally prophage

Gally is the most induced LF82 prophage *in vitro* in unstressed conditions and ciprofloxacin increases this induction even more. This raised the question of a putative negative effect of Gally on the survival of LF82 inside macrophages, an environment that provokes genotoxic stress to LF82 as judged by the SOS response level (24, 26). To test this hypothesis we compared the survival of the wild-type and ΔGally strains upon macrophage infection (Fig 6). Survival was not significantly increased with the mutant compared to the wild-type strain, neither at 6 hours nor at 24 hours post-infection, showing that the macrophage survival of LF82 is not diminished by the presence of the Gally prophage.

**Fig 6.**
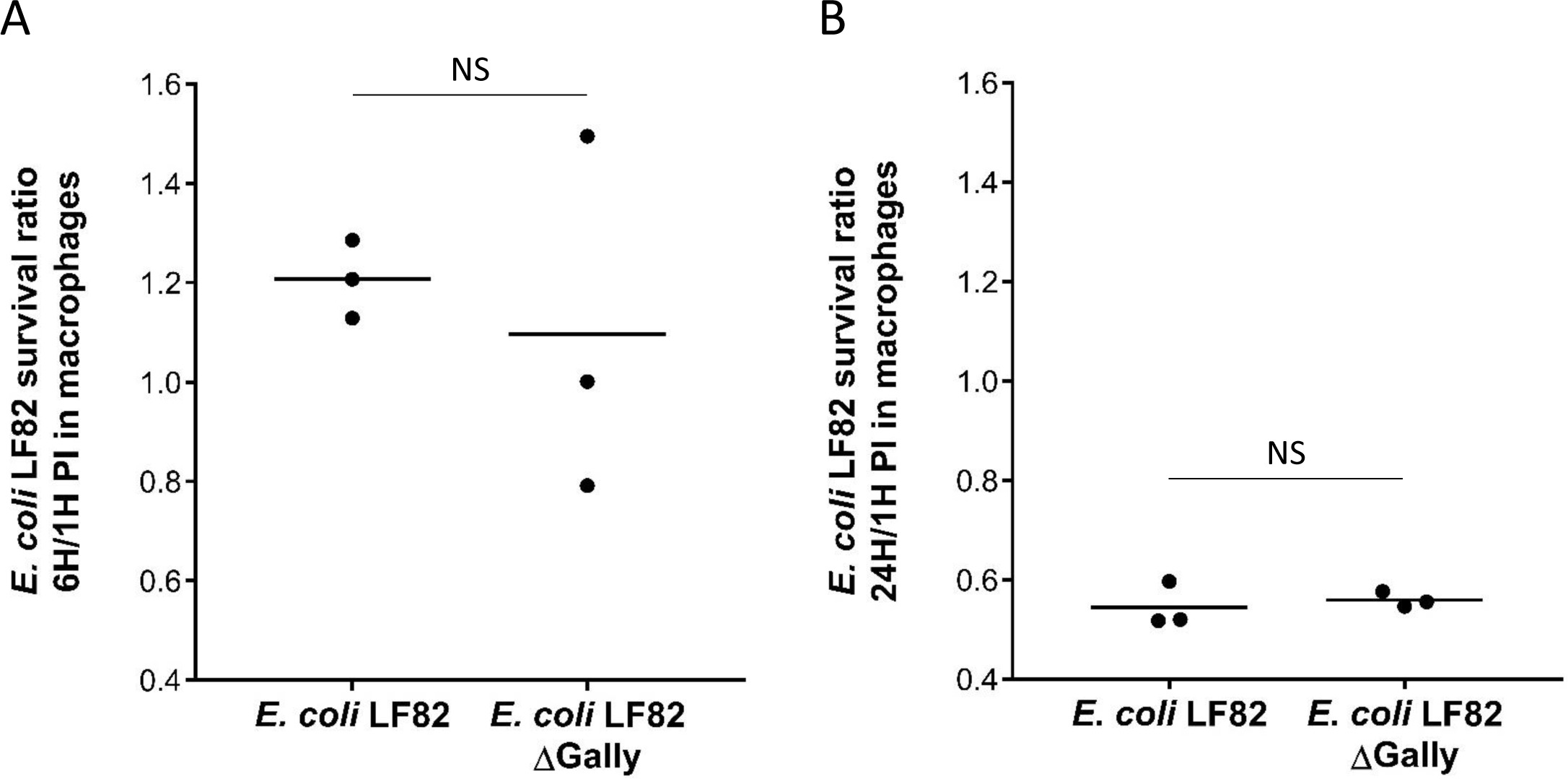
LF82 survival in THP-1 macrophages remains similar in absence or presence of the Gally prophage. *E. coli* LF82 survival (CFU/mL) after 6 (A) or 24 (B) hours in THP-1 macrophage was compared to the initial amount of endocytosed LF82 bacteria (1 hour post-infection as reference). Black dots represent values from three biological replicates obtained after independent macrophage infections and horizontal black lines represent the mean values. NS: not significant (t-test, pvalues > 0.6).

### Gally induction is prevented in macrophages

The absence of any negative effect of Gally on the survival of LF82 in macrophages strongly suggested that Gally particles were not produced in this genotoxic environment, contrary to the *in vitro* situation. We therefore investigated more precisely the fate of Gally upon macrophage infection.

Using our previously published transcriptomic analysis (26), the transcription profiles of the Gally prophage in LF82 bacteria internalized in macrophages (6 hours post-infection) or cultured *in vitro* (Fig 7, S1 Tab) were compared. The LB profile revealed high levels of the C2 repressor transcript (functional homolog of the Lambda CI), but also a background level of expression of all other genes, which might be in line with the elevated production of Gally virions. Within macrophages, the Gally region exhibited a clearly different pattern: first, the five rightmost genes of the prophage, including the Mnt repressor and a tail spike, were highly induced (7 to 50-fold). In addition, the leftward region of the prophage (replication module, from gene *c2* down to the last gene before *xis*) was transcribed 2 to 10-fold above its *in vitro* level, while several key structural genes of the rightward region, including *terS*, *terL* and *portal* as well as two genes coding for DNA injection proteins, were repressed 2 to 7-fold (Fig 7). Among the five transcriptional regulators encoded by Gally, the rightmost operon including the *mnt* repressor gene was upregulated 50-fold, C2, Cro and C1 were upregulated 4 to 7-fold and Roi and Rha was unaffected. This suggests that in LF82 bacteria growing in macrophages, an active mechanism is harnessing the transcriptional program of the prophage to prevent DNA encapsidation, and probably downstream events, including bacterial lysis while the replication and recombination module are induced.

**Fig 7.**
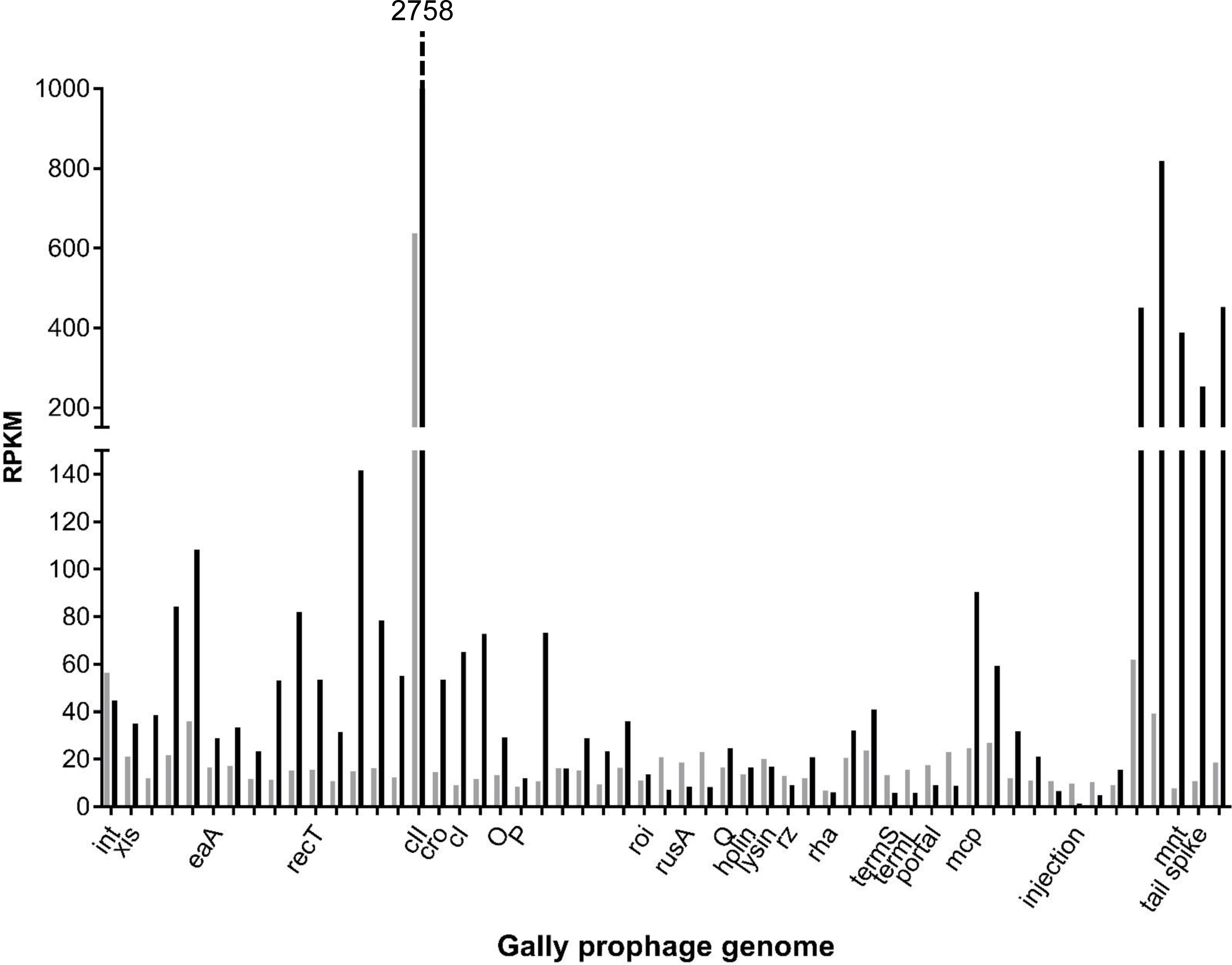
The transcriptional profile of Gally is profoundly modified in LF82 bacteria extracted from macrophages. Transcriptomes from two LF82 samples extracted from an overnight LB culture (grey bars) or from THP-1 macrophages after 6 hours of infection (black bars) were normalized for sequencing depth and gene length (RPKM, y-axis) and is presented for the 55 genes of Gally.

To further explore this possibility, we performed a qPCR estimation of the viral particle concentrations after 1, 6 and 24 hours of LF82 infection of macrophages. Concentration of Gally particles was highly variable among replicates (from 3.7x10^3^ to 2.3x10^5^ genomes/mL, Fig 8A) but remained constant over time within each replicate. In these samples, bacterial concentrations ranged between 2 to 5x10^6^ CFU/mL. Therefore, using extreme values, the virions per bacterium ratio varied between 10^-3^ to 6x10^-2^, compared to 1 *in vitro* in exponentially growing unstressed cultures. This result suggests, assuming a similar efficiency of phage recovery in both growth conditions, that Gally virions are 20 to 1000-fold less abundant per bacteria in macrophages, compared to *in vitro* during exponential growth. We also investigated whether productions of Cyrano and Perceval, which virion yield was increased by a ciprofloxacin treatment *in vitro*, were affected in macrophages. A similar trend of prophage repression was observed, with most samples exhibiting undetectable virions, and two Cyrano samples having in average 4.6x10^3^ genomes/mL (Fig 8A, 1 hour post-infection). The ratio of Cyrano virions per bacterium was therefore 9x10^-4^ in macrophages, compared to 8x10^-3^ in exponentially growing unstressed cultures.

**Fig 8.**
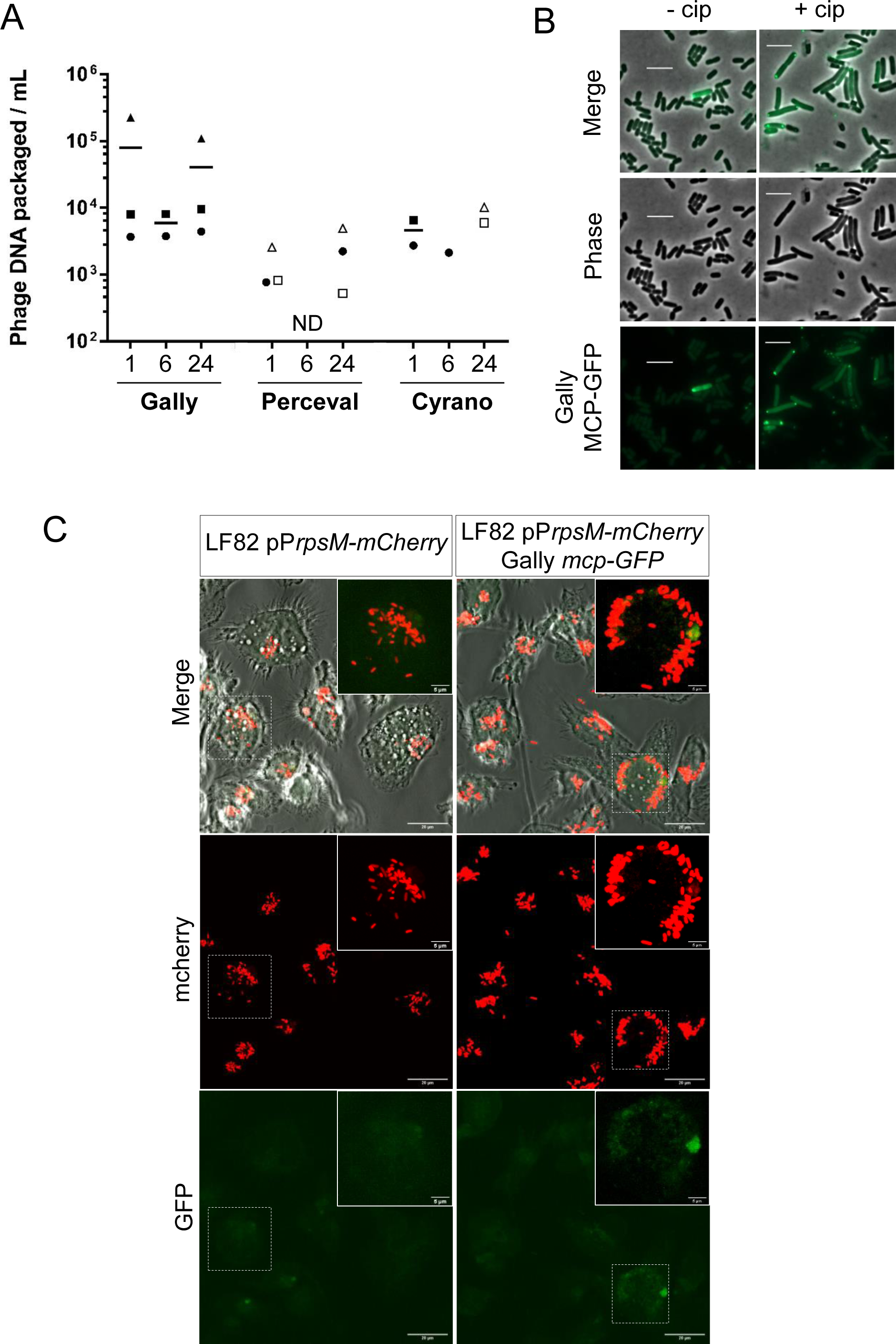
Gally particles are barely detectable in macrophages infected with LF82. A. PCR quantification of Gally, Perceval and Cyrano virions produced from LF82 bacteria after 1, 6 or 24 hours of THP-1 macrophage infection. Points of the same shape correspond to values obtained for the same biological replicate after various times of THP-1 macrophage infection (x-axis, in hours). Black symbols correspond to replicates that are greater than their associated LF82 genomic DNA contamination and are used to calculate the mean (horizontal lines), while white symbols are below and are excluded from the mean calculation. ND: not detected. B. Gally phage induction was followed by the MCP-GFP fusion production in *in vitro* ciprofloxacin-treated LF82 cultures. Here are shown snapshots of LF82 bacteria (strain MAC2606) 60 minutes after ciprofloxacin treatment (+cip) or of untreated cells (-cip). Scale bars indicate 5 µm. C. Confocal imaging of THP-1 macrophages after 6 hours of infection with the LF82-pP*rpsm-mCherry* (OEC2425) (left panel) or LF82-pP*rpsm*-*mCherry* Gally *mcp-GFP* (OEC2481) (right panel). White framed areas correspond to zooms of the white dashed framed parts. Scale bars indicate 5 or 20 µm, as indicated.

To determine the contribution of induction frequency to these net changes in phage per bacterium ratio, we measured the frequency of Gally induction by identifying cells that expressed the major capsid protein (MCP) of Gally fused to the GFP. This fusion did not allow the formation of Gally virions (as verified by qPCR, data not shown), but it permitted the visualization of fluorescent dots within intact or recently lysed bacteria. A population of 1 hour ciprofloxacin treated cells produced 44% of such fluorescent bacteria (616 fluorescently labelled cells on 1387 observed). The frequency of spontaneous Gally induction was 0.37% (20 fluorescently labelled cells on 5337 observed) (Fig 8B). Within macrophages, LF82 express the SOS response to tolerate genotoxic stress produced by the lysosomes (24, 26). Upon macrophage infection however, none of the imaged LF82 *mcp-GFP* bacteria displayed fluorescent dots, and only one lysis event was detected over a total of 2232 observed bacteria (Fig 8B, 6 hours post-infection, see S3 and S4 Fig for 40 min. and 24 hours post-infection, respectively). Gally induction frequency was therefore approximately 0.04% in LF82 infecting macrophages, around 10-fold lower than *in vitro* unstressed growth conditions.

## DISCUSSION

The five predicted prophages of strain LF82 were spontaneously produced in exponential growth phase under unstressed culture conditions. However, based on our quantification of free virions (unadsorbed at the bacterial surface), a clear gradation was observed: Gally, a P22-like podophage had the highest production level of virions (estimated ratio of 1 virion per bacterium), Cyrano and Tritos, a SSU5-like and a Lambda-like phage, respectively, produced some 100 to 150-fold less free virions than Gally. Finally, spontaneous production levels were the lowest (1300 to 2500-fold lower than Gally) for Cartapus and Perceval (estimated from the ΔGally mutant for the latter), a P2 and a Lambda-like phage respectively.

The high abundance of Gally virions in the supernatant of LF82 bacteria grown exponentially is unusual but not so exceptional. For instance, prophage BTP1 from *Salmonella typhimurium* ST313, that shares homology with P22 and HK620, is also highly spontaneously induced *in vitro*, giving rise to 10^9^ virions per mL of a stationary phase culture of the host strain (38). This high production of virions is however not a hallmark of P22-like viruses since BTP1 is ∼10,000-fold more produced than P22 in the native P22-host *Salmonella enterica* serovar Typhimurium LT2 (38). Our estimation of the spontaneous Gally induction *in vitro* (0.37%) is also similar to the one calculated for BTP1 (∼0.2%, (38)). As Gally, BTP1 is induced by a SOS promoting, genotoxic agent (38). However, in contrast to BTP1 (38), Gally virions were not able to infect their native host deleted of the Gally prophage, nor any other strains tested in this study. We are therefore unable to conclude about the infectivity of the Gally virions.

A transcriptome analysis allowed to uncover systematically the morons (i.e. genes transcribed by dormant prophages which are not necessary for its lysogenic cycle, and rather provide the host with additional functions) encoded by these prophages, and we demonstrate the presence of 30 moron genes, with more than a third of them being of unknown function. Clearly, efforts should be placed in the future to better understand the biological function of phage moron genes in natural environments. Interestingly, although the Gally prophage is highly present in *E. coli* strains associated to Crohn disease patients (22), it does not carry any moron genes in its genome.

The elevated production of Gally virions allowed the identification of its lateral transfer activity. Lateral transduction was first described for prophages of *Staphylococcus aureus* (34), and then identified as well for phage P22 (39) and phages from *Enterococcus faecalis* VE14089 (40). Whether this lateral transduction contributes to the expansion of *E. coli* strains adapted to survival in dysbiotic microbiota is unknown at present. We searched for virulence or adaptation genes in the transduced regions, and found none, except those present within prophage Perceval. Indeed, we estimated that 90% of particles containing Perceval DNA were probably due to Gally transducing particles. Prophage evolution might therefore depend in part on this lateral transduction process, whereby a region of a Lambda-like prophage could be exchanged for Gally- transduced Perceval genes. Perceval encodes two morons with known functions that are relevant for different human environments: (i) the SitABCD transporter might help LF82 bacteria to scavenge metal ions (iron or manganese) during macrophage infection and (ii) the Emr multidrug exporter might be beneficial as well for the intra-vacuolar life-style of LF82. But what Gally’s lateral transduction would bring to its LF82 host remains unclear at this time.

Bodner *et al*. demonstrated recently that a K12 lysogen has some 30-fold increased levels of prophage induction once inside macrophages compared to the induction observed *in vitro* on an agar pad (15). We show here that such phenomenon is not the case for LF82, at least for its most active prophage *in vitro*, Gally, whose particles are barely detected in macrophages, whereas they abound *in vitro.* This decreased Gally production may be due, at least in part, to the repression of its lytic cycle in LF82 infecting macrophages, since this latter is around 10-fold less induced than in *in vitro* unstressed conditions. It may be however that the lytic cycle is initiated but then stopped at a later stage. Indeed, transcriptomic data suggest that a dedicated control prevented the transcription of genes needed for phage DNA encapsidation. The Gally behavior is reminiscent to us of the phage φ10403S from the *Listeria monocytogenes* lysogen 10403S. In this strain as well, prophage induction initiates upon vacuolar invasion of *L. monocytogenes*, but then a dedicated process, controlled by the prophage transcriptional activator Llga, prevents late genes transcription (41). Whether a similar process takes place for Gally is unknown at present, but this study reveals that LF82 has evolved in order to control the lytic cycle of Gally inside macrophage, a prophage that is hyperactive in other growth conditions, rather than deleting it.

The φ10403S prophage-dependent “active lysogeny” process observed in *L. monocytogenes*, in which its excision occurs but its lytic cycle is interrupted, is associated with a genetic switch (42). Prophage φ10403S insertion splits the *comK* gene, which function is restored upon excision of the prophage, and permits vacuolar escape of *L. monocytogenes* to the macrophage cytosol, where the bacteria are able to multiply. Gally prophage is inserted between the *torT* and *torS* genes involved in the regulation of the *torCAD* operon coding for TMAO reductase (28). Prophage integration at this site is known to affect the regulation of *torS* and consequently of the *torCAD* operon (27). Moreover, we found a putative promoter in Gally, positioned at a place similar to the one characterized at the left boundary of the prophage HK022, which regulates the expression of *torS* in *E. coli* MG1655 (27). Yet, as far as we could detect, the transcription profile of the Gally prophage neighboring genes *torS* and *torT* was not affected upon LF82 macrophage infection, suggesting that Gally prophage was not excised inside macrophages. Moreover, absence of the Gally prophage did not affect the survival of LF82 inside macrophages, indicating that Gally has no role in this cellular environment. It is intriguing to see that the Gally prophage is well conserved in *E. coli* genomes associated to Crohn’s disease (22), but does not provide advantage to its AIEC host in macrophage in our infection conditions. However, the lack of a role in macrophages does not exclude a role of Gally in a different setting such as in the gut lumen or in external environments where TMAO is also available. TMAO respiration is performed by *E. coli* under anaerobic but also aerobic conditions (39). In the latter case, only a portion of the bacterial population expresses *torCAD*, leading to the proposal that TMAO respiration under aerobic conditions may facilitate bacterial adaptation to anaerobic conditions (43). The insertion of HK022, and most likely Gally in LF82, between *torT* and *torS* represses *torCAD* expression, and thus should inhibit this adaptive advantage (27). In this latter study, the authors proposed that the phage-dependent repression of *torCAD* could increase the growth rate of the bacterial host in presence of oxygen and TMAO, thus the dissemination of the phage in these growth conditions, most probably outside the gut. Remarkably, the HK022 prophage does not inhibit *torCAD* induction under anaerobic growth conditions (27), indicating that phage insertion at this site should not affect the physiology and competitiveness of a lysogenic strain in the gut. Thus, Gally could increase, when not induced, the competitiveness of LF82 under certain aerobic growth conditions. The subpopulation sacrificed by prophage induction would allow the dispersion of LF82 genes by lateral transduction to receptor strains, yet to be identified. Clearly, the intricate details of the “symbiosis” between a temperate phage and its host have novel shades that we just start uncovering.

## MATERIAL AND METHODS

### Bacterial strains

Table 2 lists the bacterial strains, and Table 3 the oligonucleotides used in this study. Unless otherwise stated, cultures were propagated in LB Lennox broth (5 g/L NaCl instead of 10 in regular LB), at 37°C under agitation. The LF82 ΔGally strain (MAC2225) was obtained by curation of the prophage using ciprofloxacin. For this, a culture of exponentially growing LF82 in LB at 37°C (OD600 = 0.2) was diluted ten-fold and treated with 2 µg/mL ciprofloxacin during 30 minutes. Cells were washed with LB and plated on LB agar plates. After incubation at 37°C, individual colonies were screened by PCR for the absence of the Gally prophage using primers JC206 and JC207 that hybridize downstream and upstream the prophage on the bacterial chromosome. Integrity of the *attB* region remaining upon excision was verified by sequencing the PCR fragment generated.

**Table 2.** Strains used in this study.

| Strain | Genetic background | Origin |
| --- | --- | --- |
| MAC2204 | *E. coli* LF82 | NC_011993, (21) |
| MAC2218 | *E. coli* LF82 $\Delta bla$ | (25) |
| MAC2225 | *E. coli* LF82 $\Delta Gally$ | This study |
| MAC2459 | *E. coli* LF82 *Gally-mcp-GFP*, KanR | This study |
| MAC2606 | *E. coli* LF82 *Gally-mcp-GFP* | This study |
| OEC2481 | *E. coli* LF82 *Gally-mcp-GFP*, pPrpsm-mCherry | This study |
| OEC2425 | *E. coli* LF82 pPrpsm-mCherry | (26) |
| MAC1403 | *E. coli* MG1655 *hsdr::KanR* | (2) |
| MAC2266 | *E. coli* C+ | ATCC8739 |
| MAC2294 | *E. coli* TD2158-PL4 (phage-free, susceptible to HK620 phage) | (46) |
| MAC2310 | *E. coli* UGB 2668 ( MG1655 *wbbl*+) ) | Pasteur Institute |
| MAC2267 | *E. coli* DH10B | (47) |

**Table 3.**
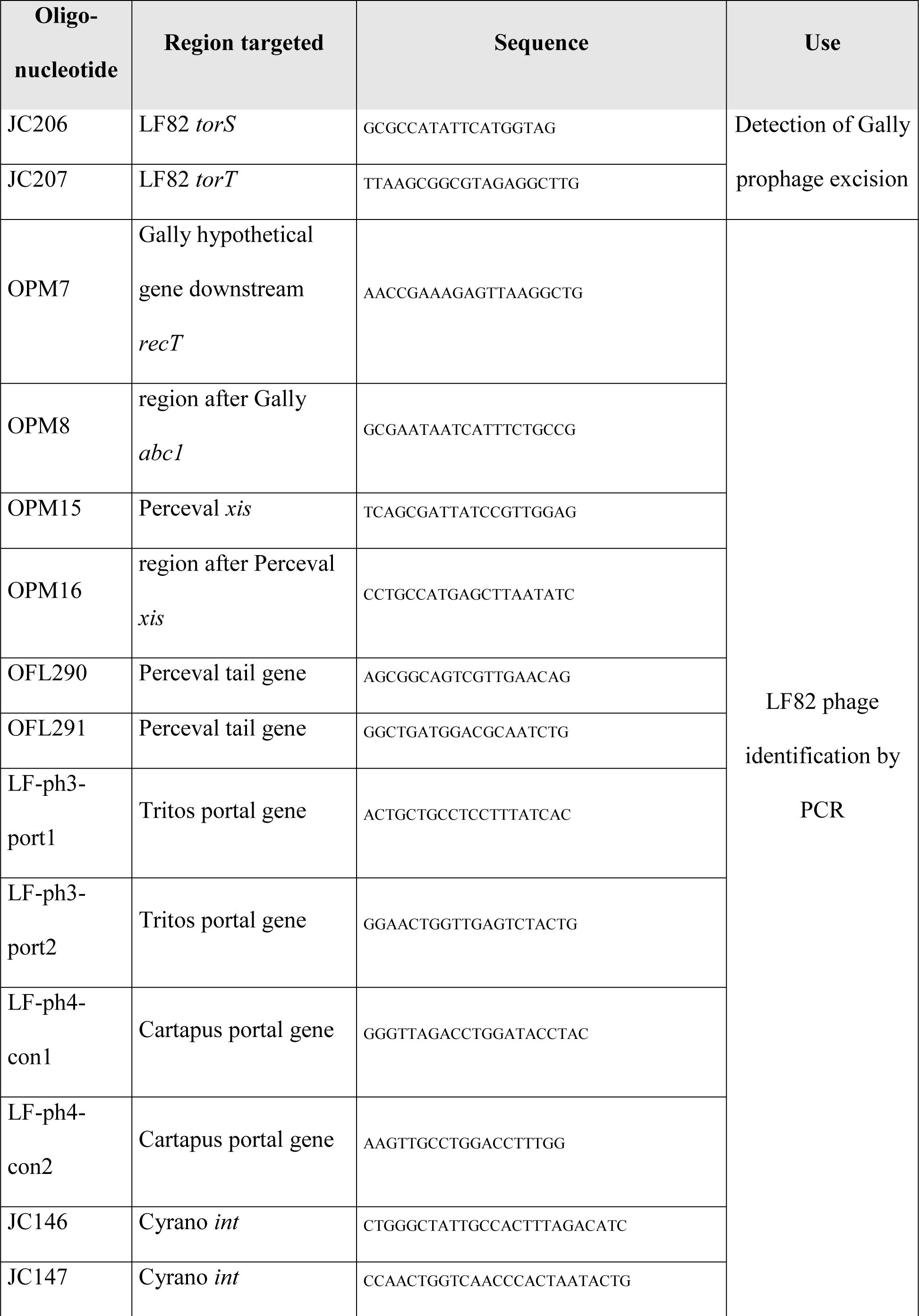
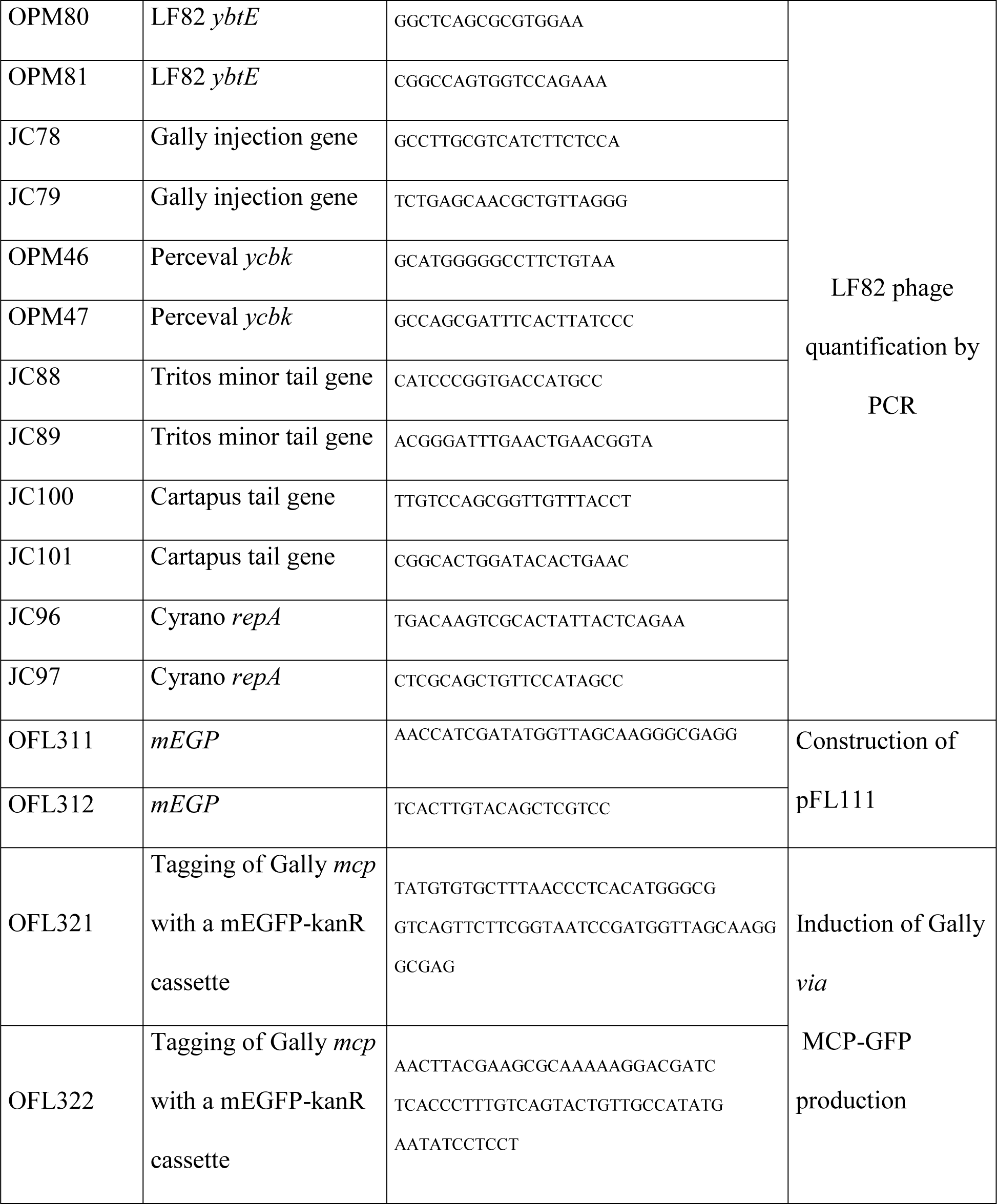
Oligonucleotides used in this study.

Strain MAC2459 (Δ*bla gally-mcp-GFP,* KanR) was obtained by integration in MAC2218 of a mEGFP-KanR cassette at the 3’ end of the *mcp* gene of the Gally prophage, by recombineering (44). In a prior step, to place *mEGFP* next to a kanamycin resistant gene, *mEGFP* was amplified with OFL311 and OFL312 from pKD3 pR:*mEGFP* (I. Matic, plasmid collection) and cloned into a ClaI/BmgBI double digested pKD4 (44). The resulting plasmid, pFL111, was then used to generate the PCR substrate for MAC2459 construction, with oligonucleotides OFL321 and OFL322. The KanR cassette was then deleted from MAC2459 *via* a Flp-FRT recombination, using the pCP20 plasmid (45), giving strain MAC2606.

Strain OEC2481 was obtained by transformation of the MAC2606 with plasmid pP*rpsm*-*mCherry* that expresses mCherry constitutively (26).

### Prophage detection and annotation

Prophage region predictions on the *E. coli* LF82 genome (NC_011993, (21)) were initiated with PHASTER (http://phaster.ca (48)), then “intact” and “questionable” regions were validated by manual inspection. Regions containing phage genes from replication, capsid, lysis and lysogeny modules were confirmed as complete prophages; we further verified the absence of genes specific of integrative plasmids or insertion sequences. To annotate hypothetical genes, a BLASTP search against the viruses taxid 10239 database was performed with default values, and the annotation of sequences producing significant alignments were transferred to the query when either experimental evidence of function or conserved domains were detected. For hypothetical proteins without BLASTP hit, sequences were analyzed for Pfam matches (49).

### Transcriptome analyses

The transcriptomic data reported in (26) were utilized. Moron detection: two ‘*in vitro*’ growth conditions, LB stationary phase, and growth for 1 hour in NMDM medium (used for macrophage cultures), were selected for moron detection (two replicates each time). Read counts, after a mapping step with Bowtie2 onto LF82 chromosome and Cyrano episome, and removal of remaining ribosomal RNA reads, were normalized in RPKM, i.e. per million reads per gene length in kb (similar to the “densities” reported in (26), except kb are replacing bp here). To detect moron in each prophage region, the “local” median value of RPKM per prophage was first computed. Genes which in all *in vitro* transcriptomes (4 samples) had an RPKM value 5-fold above this background signal due to overall repression of prophage genes, were considered morons, unless they encoded well defined phage genes (such as the master repressor, sometimes the integrase, or an anti-terminator). Comparison of Gally transcriptomic profiles: normalized reads from the LB stationary phase samples and infected macrophage samples 6 h post-infection, at which SOS response peaks, are displayed side by side.

### Homology between E. coli LF82 prophages and reference phages

Related phages were searched in the nr/nt nucleotide collection of the NCBI by BLASTn, within the Viruses taxid 10239 as of March 2020, using the megaBLAST parameters. For each prophage, the type phage of the viral species (as defined by the ICTV, ictv.online.org) and the closest phage were retained for comparison through genomic alignments. Alignments were realized using the R package GenoPlotR (50), based on tBLASTx to generate the comparison files and using a filter length of 50. Images were generated with the plot_gene_map function with the blue_red global color scheme.

### Sequencing of E. coli LF82 virome

One liter of LF82 culture grown under agitation at 37°C in LB to an OD600 ∼1 was centrifuged for 7 minutes at 5,000 g at 4°C. Supernatant was filtrated on a 0.2 µm membrane and nanoparticles were precipitated with 10% PEG 8000 and 0.5 M NaCl during an overnight incubation at 4°C. The preparation was then centrifuged for 30 minutes at 5,000 g and supernatant was removed. A second centrifugation for 5 minutes was added to completely eliminate the supernatant. The pellet was resuspended in 2 mL of SM Buffer (50 mM Tris-HCl pH 7.5, 100 mM NaCl, 10 mM MgSO4) and treated during 30 minutes at 37°C with 4 µg of RNAse and 2 U of Turbo DNAse (Ambion, Ref AM2239). Another incubation of 30 minutes at 37°C with an additional quantity of Turbo DNAse (2 U) was added to maximize the removal of bacterial DNA from the sample. Then Turbo DNAse was inactivated with 10 mM of EDTA pH 8. To extract phage DNA, we performed two phenol-chloroform–isoamyl alcohol (25:24:1) extractions followed by a chloroform-isoamyl alcohol (24:1) purification step. Then DNA was precipitated with two volumes of pure ethanol at 4°C and pelleted with a full-speed centrifugation for 5 minutes. Ethanol was eliminated by evaporation and the DNA pellet was resuspended in 40 µL of 10 mM Tris-HCl pH 8. Double-stranded DNA concentration was measured with a Qubit (dsDNA Broad range assay kit, Invitrogen, Ref Q32850) at 88 ng/µL, and 525 ng were sent to Eurofins for Illumina High seq paired-end sequencing (2 million read depth).

Reads obtained were filtered with TRIMMOMATIC (51) to keep only those of high quality using the command ILLUMINACLIP:TruSeq3-PE.fa:2:30:10 LEADING:3 TRAILING:3 SLIDINGWINDOW:4:20 MINLEN:125. Remaining reads were mapped with Bowtie2 (-N 0 -L 32) (52) on a sequence that concatenated the LF82 chromosome (CU65163, (21)) and the “LF82 plasmid”, now Cyrano, genome (CU638872, (21)). Finally the coverage information was extracted using Tablet (53) and represented with ggplot2 on R. Coverage corresponding to the mean genomic DNA contamination was calculated by using unmapped reads from a Bowtie2 alignment on a sequence concatenating the all 5 prophages (7.6 reads/bp).

To precisely delimit prophage borders, clipped reads providing evidence of the prophage recircularization were identified by Tablet identification of reads mapped on the 5’ and 3’ ends of each prophages followed by a BLASTn confirmation. These boundaries were also verified by PCR amplification and sequencing and are reported in Table 4.

**Table 4:** Prophage boundaries on LF82

| Name | start | end |
| --- | --- | --- |
| Gally | 998943 | 1037635 |
| Perceval | 1172884 | 1223241 |
| Tritos | 1576972 | 1632704 |
| Cartapus | 2734926 | 2770036 |

### Phage isolations

Perceval was isolated from the LF82 ΔGally strain (MAC2225), using a 5 mL exponentially growing culture treated with 2 µg/mL ciprofloxacin (4h30). The culture was then centrifuged for 4 minutes at 11,000 g at 4°C and filtrated on 0.2 µm membrane. As no phage plaque was observed on *E. coli* DH10B in this supernatant, an enrichment step was added: 250 µL of supernatant was adsorbed to 500 µL of DH10B overnight culture, supplemented with 10 mM MgSO4 and 1 mM CaCl2 at 37°C for 10 minutes, diluted in 50 mL of Lennox and incubated overnight at 37°C. This time, an agar overlay (10 g/L bactotryptone, 2.5 g/L NaCl, 4.5 g/L agar) with 40 µL of supernatant and 100 µL of DH10B overnight culture, plated and incubated at 37°C overnight allowed to detect small plaques, either turbid or clear. Both plaque types were streaked for purification, and large stocks were prepared by lysis confluence on plates, and recovery in SM buffer by diffusion (1 h, 4°C) followed by filtration (0.2 µm). PCR analysis using diagnostic primers for each predicted LF82 prophage (Table 3) gave positive results only with the Perceval primers OPM 15-16 and OFL 290-291, and not with the others, indicating that both purified phages were Perceval.

A Tritos phage plaque was isolated once, directly from an LF82 culture supernatant plated on strain MG1655 *hsdR-* (MAC1403). After purification and amplification, PCR analysis of this phage stock gave a positive result with Tritos primers LF-ph3-port 1 and 2 primers and not with the others, indicating that the isolated phage was Tritos.

Two Gally-Perceval hybrid phages (named Galper1 and 2) were also isolated from LF82 culture supernatant, upon plating either on LF82 ΔGally (MAC2225) or MG1655 *hsdr-* (MAC1403). A single phage plaque was obtained in each case, streaked (small and clear plaque phenotype), and amplified. To exclude PCR signal coming from LF82 DNA contamination, the crude lysate was then treated with DNAse. For this, 10 µL of phage stock was diluted 1:100 in H2O and treated with 1 U Turbo DNAse (Ambion, Ref AM2239) at 37°C for 1h30 to remove bacterial genomic DNA. DNAse was then inactivated with an incubation for 30 minutes at 95°C and the sample was analyzed by PCR with the diagnostic primers (Table 3). Results were positive both with primers OPM7-OPM8 and OFL290-OFL291, indicating that the phage isolated was composed of parts of Gally and Perceval genomes.

All conditions tested to isolate the Gally phage were unsuccessful. These included i) induction from LF82 using 1 µg/mL mitomycin C (for 2h15 and 4h20), or to 0.09 µg/mL ciprofloxacin for 2 hours. ii) Plating supernatants on various strains: LF82 ΔGally (MAC2225), *E. coli* C+ (MAC2266), *E. coli* TD2158 (MAC2294) and *E. coli wbbl+* (MAC2310). iii) Plate incubation at different temperatures: 25, 30, 37 and 42°C. iv) LF82 supernatant enrichment on TD2158 (MAC2294) or MG1655 *wbbl+* (MAC2310), as follows: 250 µL of *E. coli* LF82 filtered (0.2 µm) supernatant was adsorbed to 500 µL of overnight MAC2294 or MAC2310 cultures, with 10 mM MgSO4, 1 mM CaCl2 (10 minutes at 37°C), then diluted in 50 mL of Lennox supplemented with 10 mM MgSO4, 1 mM CaCl2 and incubated overnight at 37°C.

### Virions observation by electron microscopy

1 mL of purified stocks of Tritos (1.4x10^7^ PFU/mL), Perceval (6.2x10^10^ PFU/mL) and Gally-Perceval hybrid (10^11^ PFU/mL) crude lysates were concentrated for TEM observation by successive washes in ammonium acetate following the protocol from Nicolas Ginet (CNRS, France, personal communication). After centrifugation for 1 hour at 20,000 g, 4°C, the pellets were resuspended in 1 mL 0.1 M ammonium acetate pH 7 (previously filtrated on 0.2 µm membrane). Tubes were centrifuged once again and pellets were resuspended in 50 µL 0.1 M ammonium acetate pH 7.

For Gally imaging, 200 mL of LF82 overnight culture were centrifuged for 7 minutes at 5,200 g. Supernatant was filtrated on 0.2 µm membrane and centrifuged for 3h at 143,000 g, 4°C to concentrate the virions. Resulting pellet was resuspended in 12 mL of 0.1 M ammonium acetate pH 7 before being centrifuged once again for 2 hours at 154,000 g, 4°C. The final pellet was resuspended in 30 µL of 0.1 M ammonium acetate pH 7.

Cyrano was visualized by TEM using the same protocol as the one used for Gally observation except that the LF82 culture in exponential growth phase (OD600 ∼0.3) was treated for 2 hours with 0.09 µg/mL ciprofloxacin.

Ten µL of each virion preparations were absorbed onto a carbon film membrane placed on a 300- mesh copper grid and stained with 1% uranyl acetate dissolved in distilled water. After drying at room temperature, grids were observed with Hitachi HT 7700 electron microscope at 80 kV (Elexience – France) and images were acquired with a charge coupled device camera (AMT). Finally, tails and capsids were measured using ImageJ software (54).

### Gally-Perceval hybrid genome assembly

Galper1 was entirely sequenced following the same first steps described above for virome sequencing. After read cleaning, a dereplication step was computed, using the USEARCH9 command line -fastx_uniques (55), pairs were reconstituted using FASTQ_PAIR and reads were assembled with SPADES (--careful -k 21,33,55,77,99,127 option, (56)). A single contig of 44,690 bp, corresponding to the complete genome of Galper1, was obtained (see map and virion, S2 Fig).

To test whether the recombination junctions were placed similarly in Galper2, the two regions were PCR amplified with primers OPM 50-51 and OPM52-53, and sequenced.

### Determination of minimal inhibitory concentrations of antibiotics for LF82

We first determined the ratio between OD600 and CFU/mL for LF82 (MAC2218). OD600 from three independent 18 hours cultures of LF82 were measured, and samples were plated and incubated overnight at 37°C. Colony counts indicated that a saturated LF82 culture contains about 9.6x10^8^ bacteria/mL per OD unit.

Taking this ratio into account, we then determined the minimal inhibitory concentrations (MIC) of LF82 for gentamicin (Sigma, ref G1264-1G, resuspended in H2O), cefotaxime (Sigma, ref 219380, resuspended in H2O), trimethoprim (Sigma, ref T7883-5G, resuspended in 100% DMSO) and ciprofloxacin (Sigma, ref 17850-5G-F, resuspended in 100 mM HCl) in Lennox medium by following the protocol from Wiegand & al. (57). Three independent 18 hours cultures of LF82 were diluted to 10^6^ CFU/mL and 1 mL of each was added to 1 mL of Lennox containing increasing concentrations of antibiotics (two-fold steps): 6.3x10^-2^ to 125 µg/mL for gentamicin, 9.4x10^-4^ to 9.6x10^-1^ µg/mL for cefotaxime, 2.3x10^-3^ to 2.4x10^-1^ µg/mL for ciprofloxacin and 7.8x10^-3^ to 8 µg/mL for trimethoprim. To verify the input bacterial titer, cultures containing no antibiotics were numerated on Lennox agar and incubated overnight at 37°C. Cultures were incubated at 37°C for 20 hours under agitation. MICs corresponded to the minimal concentrations of antibiotics that completely inhibits growth (OD600 below 0.05) of LF82: 15.63 µg/mL for gentamicin, 0.72 µg/mL for cefotaxime, 0.09 µg/mL for ciprofloxacin and 0.33 µg/mL for trimethoprim.

To confirm these results for microplate cultures, 50 µL of antibiotics dilutions were added to 50 µL of diluted LF82 (MAC2218) overnight culture (10^6^ CFU/mL) and plated in 96-wells plate which was closed with a semi-permeable filter (Gas permeable film, 4titude, Ref 4ti-0516/96) to prevent evaporation. The plate was incubated for 20 hours at 37°C in a Tecan fluorimeter. Using the TECAN I-CONTROL software, OD610 of each well was measured every 3 minutes after 15 seconds of orbital shaking of the plate at 158.9 rpm and a wait time of 5 seconds. 10 µL from cultures without any antibiotics were collected as previously to verify the input bacterial titer. MICs obtained from microplate cultures were close to those obtained from the test tube assay: 15.63 µg/mL for gentamicin, 0.24 µg/mL for cefotaxime, 0.06 µg/mL for ciprofloxacin and 0.125 µg/mL for trimethoprim.

### Quantitative PCR of LF82 phage concentration in uninduced and induced in vitro conditions

Overnight culture of LF82 (MAC2218) was diluted 1:500 and grown at 37°C to an OD600 between 0.2 and 0.3. Cultures were then diluted 1:2 with or without antibiotics at the MIC, and 200 µL were incubated in a 96-wells plate covered with a semi-permeable membrane for 2 hours at 37°C in a Tecan fluorimeter, until cultures without antibiotics reached a plate-reader OD610 between 0.25 and 0.3. To recover phage supernatants, the plate was centrifuged at 5,200 g for 7 minutes at 4°C and supernatants were filtrated on 0.2 µm membrane. Samples (107 µL) with similar growth profiles and final ODs were treated during 1 hour at 37°C with 2 U Turbo DNAse to remove the bacterial DNA. An incubation at 95°C for 30 minutes inactivated the DNAse and released the phage DNA from virion capsids. The final samples were diluted 1:50 and 1:100 in pure water, and 6 µL were used for the PCR quantification.

The bacterial DNA of LF82 was used as a reference point for qPCR measurements of phage copy number (1 prophage copy per genome). Genomic DNA was extracted from an overnight culture lysate treated twice with phenol-chloroform–isoamyl alcohol (25:24:1), followed by four chloroform-isoamyl alcohol (24:1) extractions. DNA was ethanol precipitated and resuspended in 10 mM Tris-HCl pH 8. DNA was quantified with Qubit (dsDNA Broad range assay kit, Invitrogen, Ref Q32850) and serial diluted in 10 mM Tris-HCl pH 8 to obtain a range from about 50 to 5.10^5^ copies of *E. coli* LF82 genome per 6 µL.

To quantify the packaged phage DNA, we used the Luna® Universal qPCR Master mix from NEB (Ref M3003E) with primers described in Table 3 at 250 nM each to target specifically each phage genome or OPM 80-81 primers to target the yersiniabactin biosynthesis salycil-AMP ligase protein encoding gene (*ybtE*) from LF82 and evaluate the bacterial DNA contamination of our samples. Nine µL of this mix were added either to 6 µL of diluted viral samples, 6 µL of LF82 genome for the qPCR reference, or 6 µL of H2O (negative control) and run in a StepOne™ Real- Time PCR System (ThermoFisher scientific) with the following program : 95°C 1 min, (95°C, 15s; 60°C, 30s) 40 cycles, followed by melting curves. Results obtained were analyzed using the StepOne Software 2.3.

### E. coli LF82 survival in macrophage

THP1 (ATCC TIB-202) monocytes (4.75x10^5^ cells/mL) were differentiated into macrophages in phorbol 12-myristate 13-acetate (PMA, 20 ng/mL). *E. coli* LF82 (MAC2204) and *E. coli* ΔGally (MAC2225) were used to infect 4.75x10^5^ THP1 macrophages. After 1, 6 or 24 hours Post-Infection (P.I), THP1 macrophages were lysed with 500 µL of 1% Triton-PBS. Lysate was plated on Lennox medium and incubated at 37°C. Colonies were counted to determine the CFU/mL after macrophage infection of each LF82 strain at each time point.

### Analyses of the Gally phage induction in vitro by epifluorescence microscopy

An overnight culture in Lennox medium at 37°C of the MAC2606 strain was diluted 100-fold in fresh medium. At OD600 ∼0.3, 0.09 µg/mL of ciprofloxacin was added. At 0 and 1 hour after ciprofloxacin addition, cells were deposited on slides covered with 1.5% agarose in M9 minimal medium. Cover slips were positioned and slides were examined and analyzed with a fluorescent microscope Carl Zeiss AxioObserver.Z1. Images were acquired with a 100x oil immersion objective and processed with Zen (Carl Zeiss) or Image J softwares.

### Gally phage induction in macrophage, followed by epifluorescence microscopy and qPCR

Strains OEC2481 and OEC2425 inside macrophages were observed as follows: OEC2481 and OEC2425 strains were inoculated in Lennox medium at 37°C at 180 rpm. The overnight bacterial culture was diluted 100-fold in fresh medium. Once OD600∼0.5 was obtained, macrophages THP1 (see above the monocytes differentiation protocol and infection) were infected and incubated at 37°C, 5% CO2 as described (58). After 40 min, 6 hours and 24 hours P.I, macrophages were fixed with formaldehyde 3.7% (Ref : F8775 Sigma-Aldrich) for 30 minutes at room temperature, and washed twice with PBS. Then the lamella was mounted with Dako. Imaging was performed on an inverted Zeiss Axio Imager with a spinning disk CSU W1 (Yokogawa) at 63X magnification. Metamorph Software (Universal Imaging) was used to collect the data.

The production of Gally, Perceval and Cyrano phages in macrophage was quantified by qPCR as follows: 1.2 to 1.7x10^7^ macrophages THPI were infected with the MAC2204 strain as described above. After 1, 6 and 24 hours P.I, macrophages were lysed with Triton 0.075% for 10 min at room temperature. Lysed macrophages were then scraped from the culture well and filtered on a PES-membrane of 0.2 µm. Viral particles from the samples (4 mL) were then concentrated 10-fold with 10% PEG 8000 and 0.5 M NaCl precipitation (see virome sequencing section), treated with 2 U of Turbo DNAse each, diluted in pure water and quantified with the Luna® Universal qPCR Master mix from NEB as previously described.

### Genome and reads submissions

The re-annotated genomes of the phages are available from the European Nucleotide Archive browser (http://www.ebi.ac.uk/ena/browser/view) with the following accession numbers: OV696608 for Gally, OV696612 for Perceval, OV696610 for Tritos, OV696611 for Cartapus and OV696614 for Cyrano. Raw data obtained from the sequencing of the virome will be deposited only after acceptance of the manuscript.

## ACKNOWLEDGMENTS

This work was supported by the Agence Nationale de la Recherche [Persist3R contract]. We are grateful to Christine Longin (MIMA2 platform) for her help with the TEM observations, to Alice Eon-Bertho for technical help, to Julien Lossouarn for the phage genomes and reads submissions and to the Migale platform (INRAE) for the bio-informatics environment.

## AUTHOR CONTRIBUTIONS

PM, EB, JC, GD and FL performed the experiments. PM, MDP and MAP performed bio- informatics analyses. PM, MDP, OE, MAP and FL designed and interpreted the experiments. MAP and FL wrote the manuscript.

## CONFLICT OF INTEREST STATEMENT

The authors declare that the research was conducted in the absence of any commercial or financial relationships that could be construed as a potential conflict of interest.

## SUPPORTING INFORMATION CAPTIONS

**S1 Fig.**
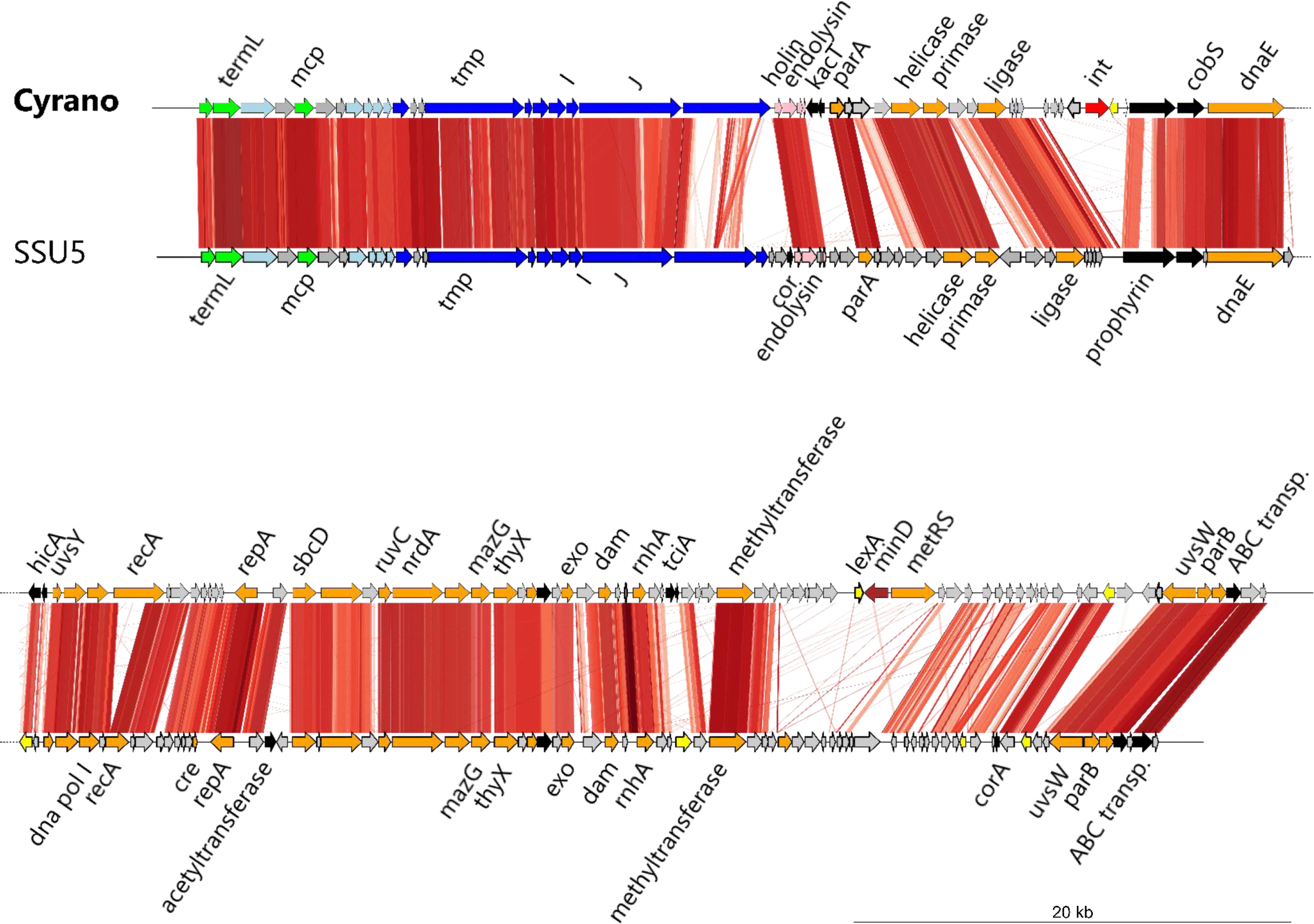
Whole genome comparison of the LF82 phage-plasmid Cyrano and SSU5. A tBLASTx comparison was performed and visualized with the R package Genoplot.

**S2 Fig.**
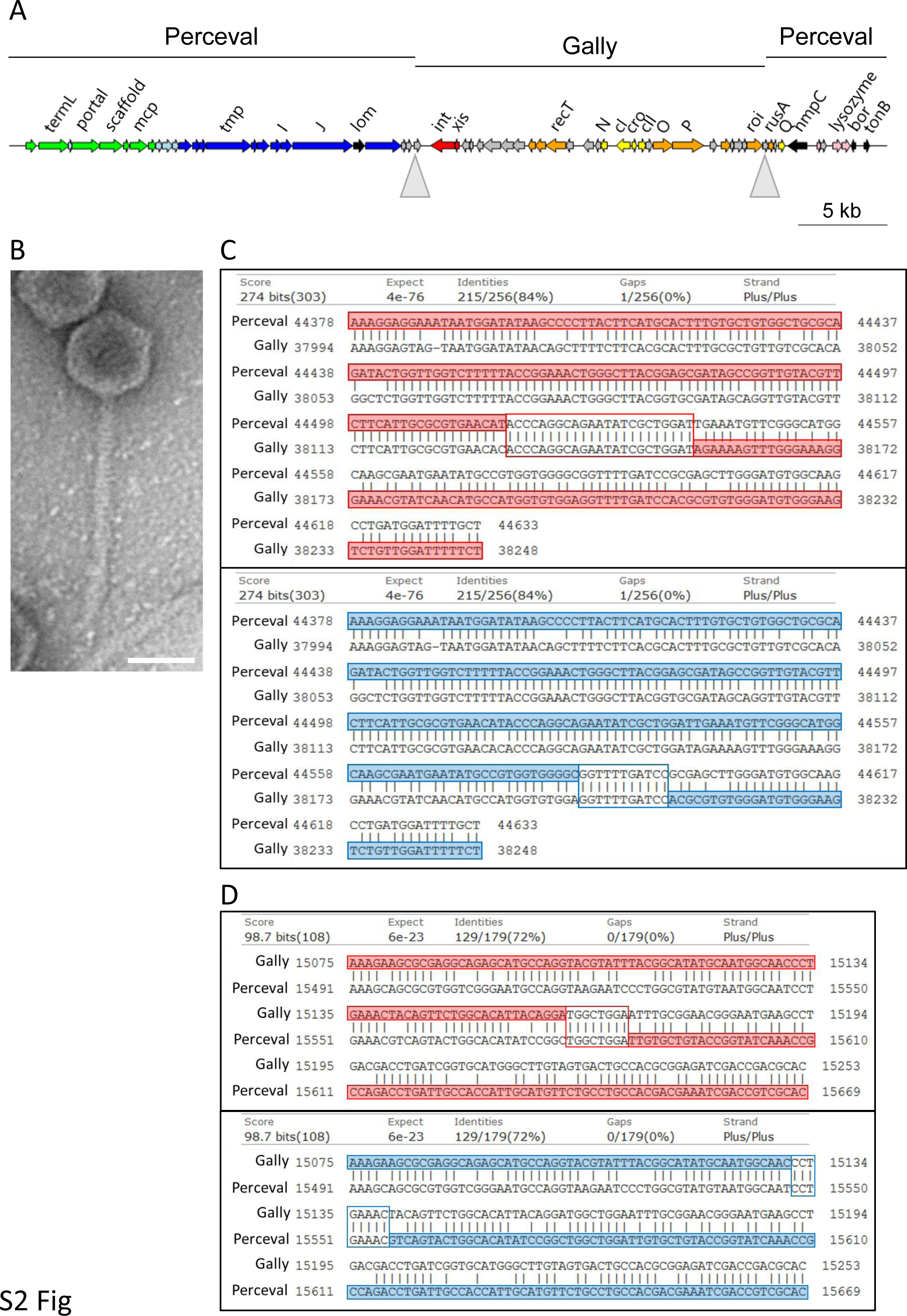
Genetic analysis of two Gally-Perceval hybrids isolated in this study. A. Genetic map of the Galper hybrids. Grey triangles indicate the two recombination endpoints between Gally and Perceval. B. Transmission electron microscopy photograph of the purified Galper1. C. Sequence analysis of the first recombination endpoint in Galper1 (upper panel, red) and Galper2 (bottom panel, blue), which occurs in a 256 bp region of partial homology between Perceval and Gally (84% identity). Sequences framed in red or blue correspond to the site where the Gally-Perceval recombination occurred. Sequences highlighted in red or blue correspond to Galper1 or Galper2 sequences in this region, respectively. D. As above, sequence analysis of the second recombination endpoint in Galper1 and Galper2, which occurs in a 179 bp region of partial homology between Perceval and Gally (72% identity).

**S3 Fig.**
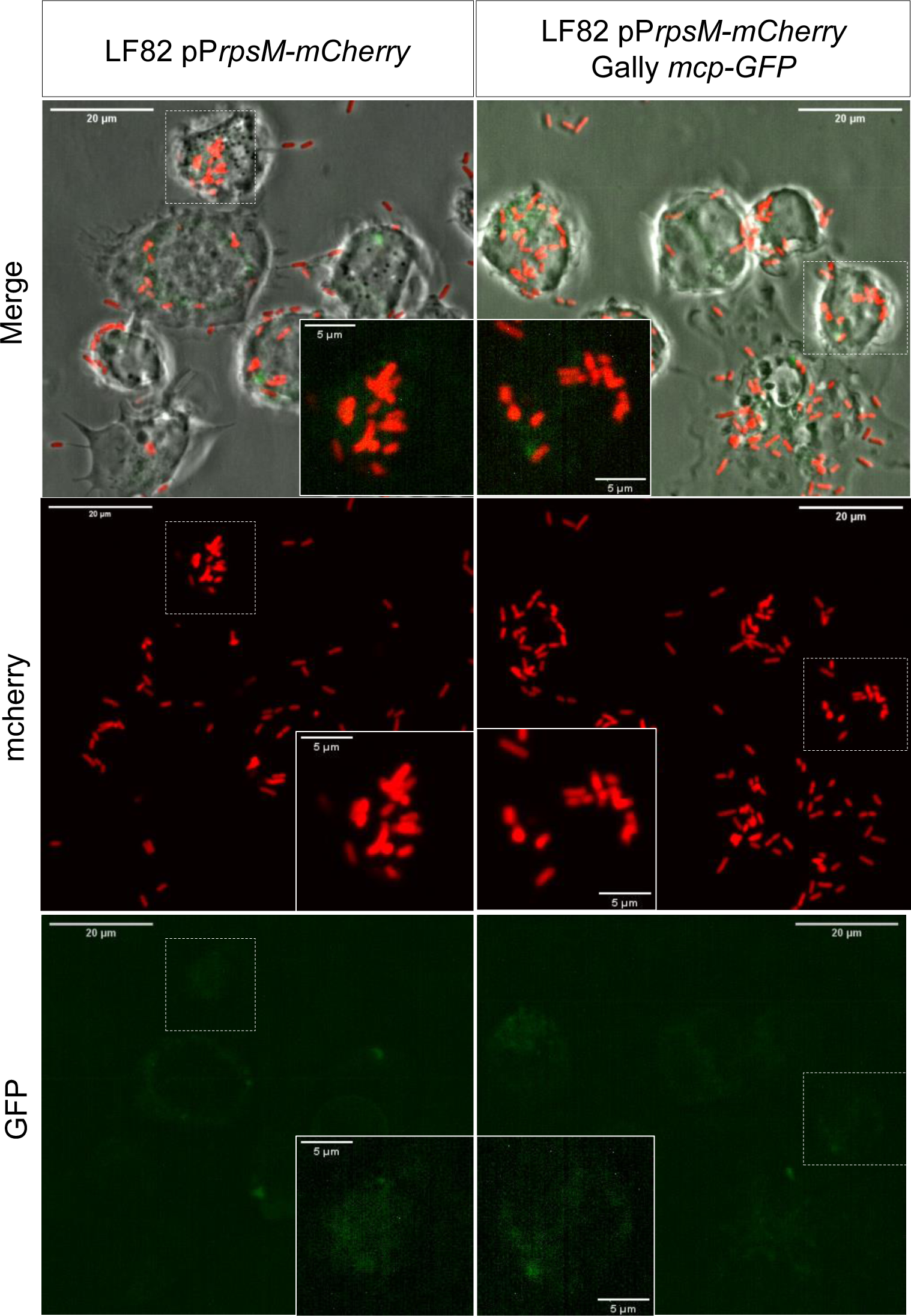
Confocal imaging of THP-1 macrophages after 40 minutes of infection with LF82- pP*rpsm*-*mCherry* (OEC2425) (left panel) and LF82-pP*rpsm*-*mCherry* Gally *mcp-GFP* (OEC2481) (right panel).

**S4 Fig.**
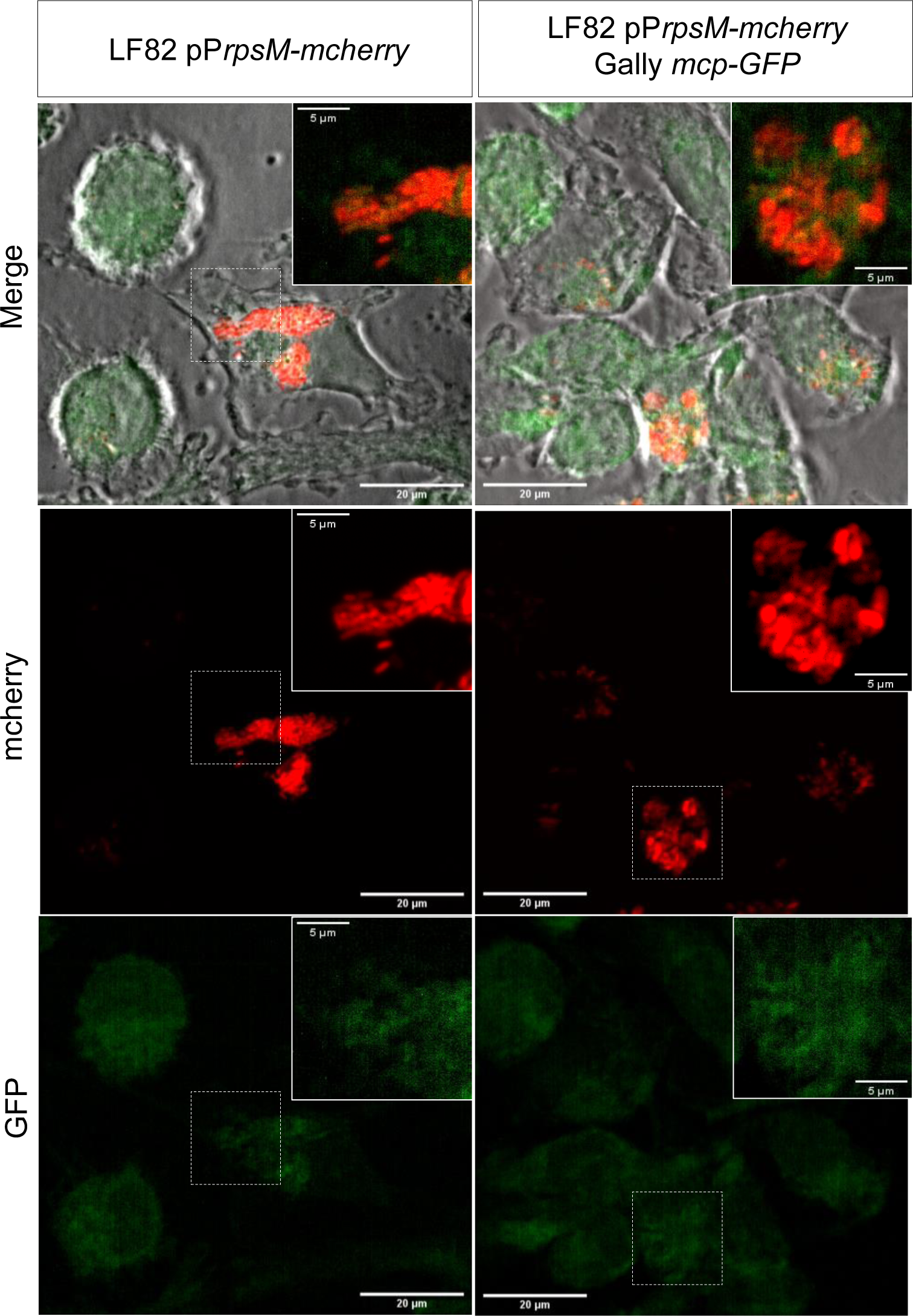
Confocal imaging of THP-1 macrophages after 24 hours of infection with LF82-pP*rpsm*- *mCherry* (OEC2425) (left panel) and LF82-pP*rpsm-mCherry* Gally *mcp-GFP* (OEC2481) (right panel).

**S1 Tab.**
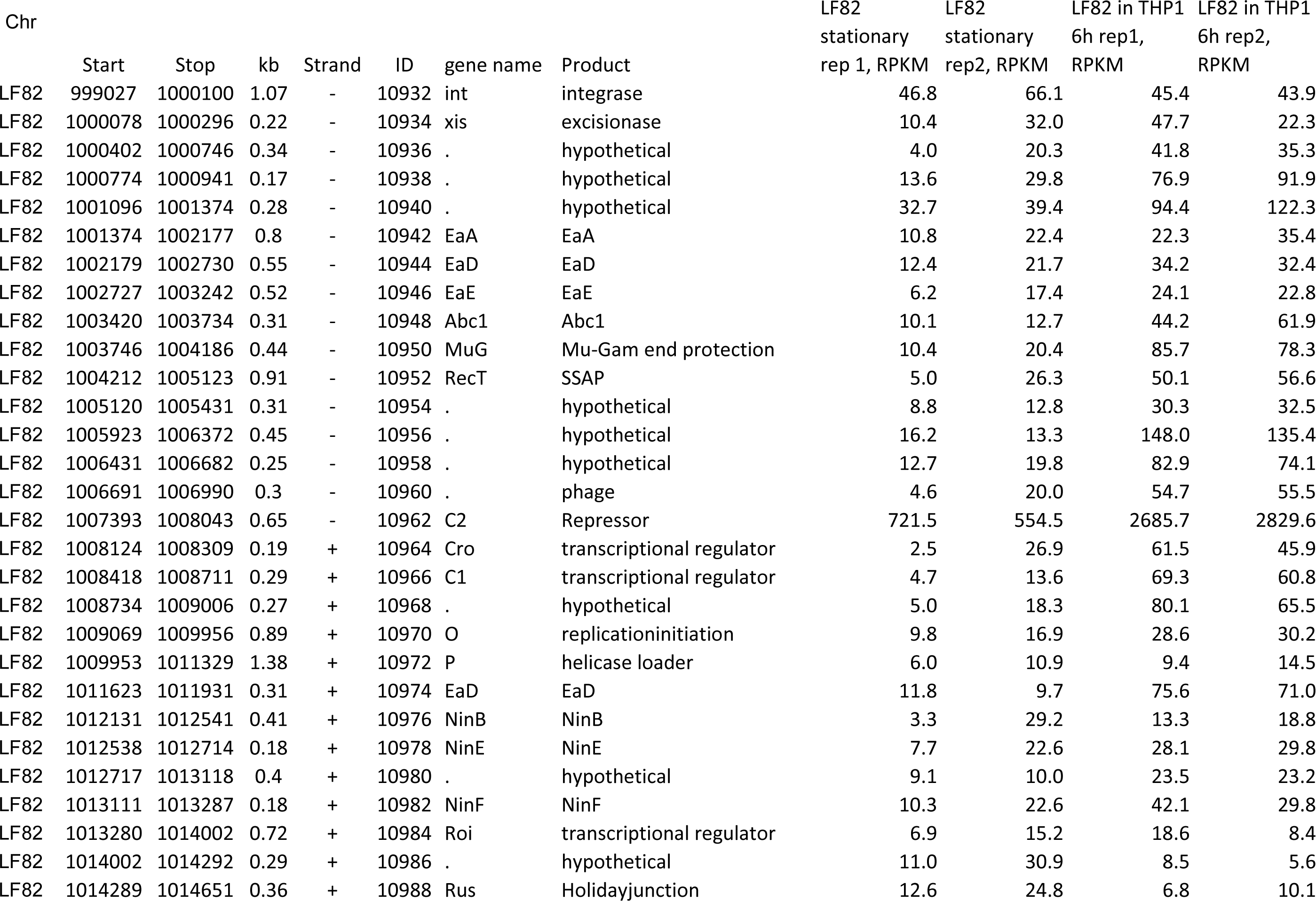
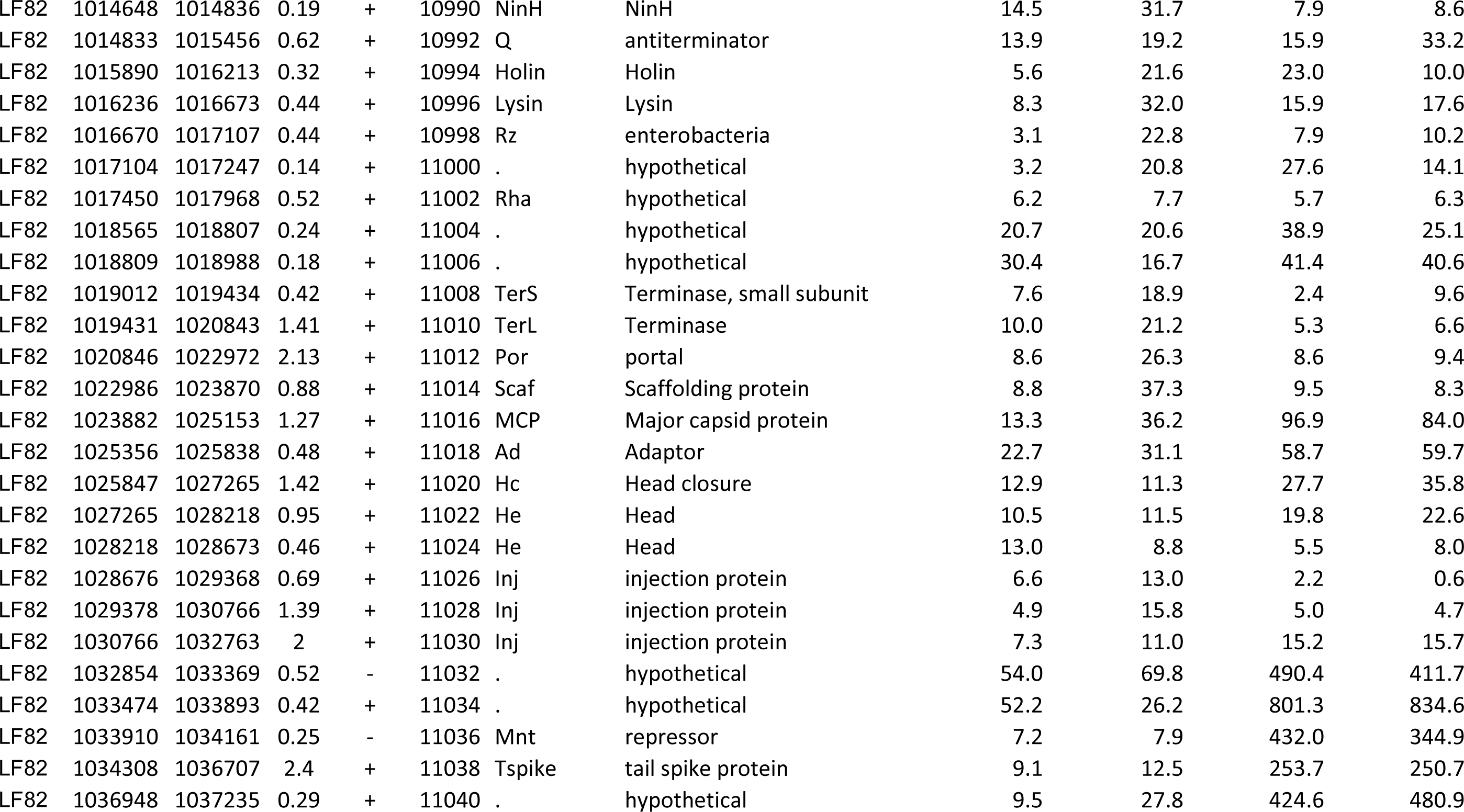
Transcriptomic analysis of Gally in stationary phase LF82 bacteria grown in LB or 6 hours post-infection in macrophage

## Notes

### Competing Interest Statement

The authors have declared no competing interest.

